# Do LLM Proposers Improve Budgeted Search for Ionizable Lipids?

**DOI:** 10.64898/2026.10.06.756984

**Authors:** Harshit Singh, Rajeev Kumar Singh, Chundru Sharath Krishna, Satya Pratik Srivastava, Rohan Gorantla

**Affiliations:** Department of Computer Science and Engineering, Shiv Nadar University, Delhi-NCR, India; School of Informatics, University of Edinburgh, Edinburgh EH8 9AB, U.K; EaStCHEM School of Chemistry, University of Edinburgh, Edinburgh EH9 3FJ, U.K

## Abstract

Efficient lipid design under limited experimental budgets is a central challenge in computational chemistry and drug discovery. We introduce LEAF (Lipid Evolutionary Adaptive Framework), a matched-budget protocol for testing whether language-model proposers improve combinatorial molecular search relative to classical baselines. The study uses a 30,044-candidate ionizable-lipid space, a frozen Gaussian-process oracle fitted to experimental potency labels, and a separate active-learning surrogate. Four language models evaluated over eight seeds and three splits show no significant improvement over genetic-algorithm or random-search baselines. For Llama 8B, 79.1% of acquisition-selected queries repeat previously evaluated archive members. Removing duplicates reduces the performance gap on some splits but increases proposal cost and can prevent budget exhaustion. Larger models reduce duplication without establishing superiority or equivalence. Exploratory scored prompts and stricter novelty controls yield positive median differences against the genetic algorithm, yet wide intervals and shared-seed dependence preclude stronger conclusions. A validated single-assay reanalysis preserves the chemotype-cold deficit at a smaller numerical scale, and exploratory expected hypervolume improvement does not reverse the result. Follow-up checks address whether proposers genuinely use injected evidence. These checks include a zero-shot leakage probe, rank-only and chain-of-thought-rationale prompts, a score-shuffle perturbation, an evidence-conflict probe, and a genuine reasoning-mode proposer. They show measurable engagement with supplied information without a corresponding performance gain. LEAF isolates proposal quality as the experimental variable and documents a concrete failure mode of zero-shot LLM proposers under controlled budgets.

## 1 Introduction and related work

Ionizable lipids influence lipid nanoparticle (LNP) delivery efficiency, while established synthetic routes support combinatorial variation of head groups, linkers, and tails (Chen et al., 2023). Experimental evaluation is expensive, so practical design reduces to a budgeted search problem in which only a small number of candidates can be tested (Settles, 2012; Gorantla et al., 2024); the method used to propose those candidates directly affects the quality of the lipids that are found. We introduce LEAF (Lipid Evolutionary Adaptive Framework), a matched-budget protocol that isolates the contribution of language-model proposers under identical evaluation oracles, acquisition functions, and query budgets. This design isolates proposal quality from evaluation and selection. Well-characterized combinatorial objectives offer limited scope for long-horizon reasoning or heterogeneous evidence integration, potential benefits under sparse supervision, literature-grounded objectives, or multistep synthesis remain untested.

Ugi three-component-reaction libraries and AGILE’s supervised screening pipeline provide established approaches (Chen et al., 2023; Xu et al., 2024), alongside synthesizable-lipid generative modeling (Ou et al., 2024). LipoAgent coordinates fine-tuned TxGemma and Qwen prediction and verification agents, including toxicity-conditioned efficiency prediction rather than budget matched search (Li et al., 2026). LUMI-lab integrates foundation models with robotic active learning, screens in vitro, and validates candidates in vivo (Xu et al., 2026), but does not compare proposal mechanisms under identical budgets. ProtAgents, protein-design swarms, and AI co-scientist study collaborative biomolecular reasoning (Ghafarollahi and Buehler, 2024; Wang et al., 2025; Gottweis et al., 2026). ChemCrow and Coscientist demonstrate chemistry-tool use (Bran et al., 2024; Boiko et al., 2023), ToolMol motivates tool-scored crossover and mutation (Zhou et al., 2026). LEAF combines this proposal pattern with Gaussian-process (GP) active learning and a continuous Tanimoto kernel (Gessner et al., 2024; Srinivas et al., 2009; Rasmussen, 2006; Settles, 2012). Generated lipids require a generalizing evaluation oracle distinct from the acquisition surrogate. Batch expected hypervolume improvement (EHVI), conditional generative multi-objective Bayesian optimization, and LLM-derived evolutionary operators within multi-objective evolutionary algorithms (Zhou et al., 2026) are complementary approaches not evaluated here. Related studies benchmark surrogate models, acquisition policies, and batch sizes for ligand screening (Gorantla et al., 2024), combine GP active learning with fragment-level SHAP explanations (Srivastava et al., 2026a), and use assay-supervised graph representation learning for molecular properties (Srivastava et al., 2026b). For protein interactions, fine-tuned protein language models support explainable affinity prediction (Singh et al., 2026b), while chain-aware modeling represents antibody heavy chains, light chains, and antigens separately (Singh et al., 2026a). These studies inform surrogate evaluation and interpretability but do not test LLM lipid proposals under matched oracle budgets. A concurrent diagnostic framework for biological reasoning models introduces perturbation and evidence-conflict tests to separate genuine input use from pattern completion (Fang et al., 2026). Section 3.4 adapts both tests to the proposer setting.

LEAF provides a matched-budget protocol with a pre-committed interpretation of null results. It evaluates four language-model proposers against genetic-algorithm, random-search, and bandit baselines, quantifies duplicate-driven query waste, tests prompt and novelty interventions, and audits implementation errors that can generate false positives. Secondary components produced no usable signal under the conditions examined. Any interpretability benefit remains qualitative rather than experimentally demonstrated.

## 2 Method and experimental protocol

### 2.1 Search space, objectives, and oracles

The general representation *x* = (*h, ℓ, t*_1_*, t*_2_*, u, b*) *∈ X* indexes a head group, linker, two tails, unsaturation, and halogenation. The implemented Ugi assembler instead uses amine, aldehyde, and isocyanide slots (*A, B, C*) and Morgan fingerprints. The general specification is

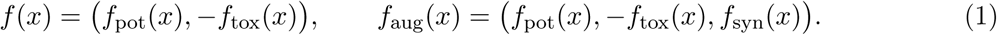

The synthesizability axis requires non-negligible variance, nearly constant route validity cannot meaningfully enlarge a frontier. Tropism heuristics are secondary filters, not objectives or evidence of organ selectivity.

#### Implemented objectives

All two-objective experiments maximize the correlated AGILE HeLa and RAW264.7 transfection-potency readouts, with both signs +1. This specification covers the primary sweep, conventional baselines, bandit, prompt variants, de-duplication, hosted models, and EHVI. Each GP head uses internal target normalization fitted on its split’s training data, the fixed hypervolume reference is (*−*5.0*, −*5.0). A deterministic toxicophore rule screens the Verifier pool only. The HeLa-only chemotype-cold reanalysis is the explicit exception: it maximizes one readout with reference (*−*5.0). Thus reported hypervolume measures potency, not a trade-off involving toxicity or synthesizability. Comparisons require matched budgets and objective definitions, a common reference alone does not remove differences in split-specific oracle fits or normalization.

The frozen evaluation oracle *O*_GT_ represents laboratory feedback in retrospective simulation. It is fitted only on each split’s training partition and must extrapolate to assembled lipids absent from the labeled library. A full-data fit would erase the intended split distinction. The separate active-learning surrogate *O*_AL_ uses the same GP architecture but is refitted each round on acquired labels. Its posterior drives acquisition, attribution, and verification without query charges, generating its own training targets would add no information. This remains model-on-model evaluation.

### 2.2 Proposal, acquisition, and secondary operators

At round *t*, the proposer receives Pareto archive *A_t−_*_1_, the reagent vocabulary, and Hypothesis Ledger *H_t−_*_1_. Each schema-constrained call returns one crossover or single-fragment mutation candidate. Exact vocabulary checks and rejection rates distinguish model output from fallback. The surrogate scores pool *P_t_* without oracle charges, after verification, scalarized multi-objective upper confidence bound (UCB), with *β* = 2, selects up to *q* candidates. All methods share this acquisition and query accounting. Random search denotes uniform random proposals, not unselected queries. The design permits *m* = *|P_t_| ≫ q*, primary experiments use *m* = 10, *q* = 5. Appendix D specifies operators and duplicate handling, Appendix E gives prompts and inference settings.

The proposed Verifier aggregates independently prompted toxicologists from at least two base models using split conformal prediction (Angelopoulos and Bates, 2023). Intervals crossing the decision threshold trigger abstention. Adaptive sampling motivates round-wise weighted or Mondrian recalibration and empirical coverage monitoring rather than guaranteed exchangeability under distribution shift (Tibshirani et al., 2019; Vovk, 2013); agreement above 95% flags insufficient diversity. No AGILE toxicity assay supports a learned head in the reported runs. Calibration scores equal deterministic RDKit labels, yielding zero-width intervals and identical decisions. These runs do not exercise epistemic debate.

Every *k* rounds, literature passages and Integrated Gradients or KernelSHAP attributions motivate structure-property hypotheses (Sundararajan et al., 2017; Lundberg and Lee, 2017). Validation requires a minimum number of matched counterfactual pairs and a paired Wilcoxon test, with every pair query charged to *B* through *O*_GT_. A bare-LLM leakage probe must precede any claimed rediscovery of the brominated-tail potency association (Xu et al., 2026); detected prior knowledge permits only a consistency check. No halogenated reagents occur in the real vocabulary, so this test was not run. The Ledger remained empty in the reported sweep. Secondary components are therefore specified architectural extensions, not empirically active mechanisms, Appendix A gives their full scope.

### 2.3 Data, controls, and statistical analysis

AGILE labels cover a complete 20 *×* 12 *×* 5 = 1,200-lipid factorial design(Xu et al., 2024). Inverting its 12,276-candidate virtual library recovers all reagent triples and adds 9 amines, 62 aldehydes, and 9 isocyanides, yielding 29 *×* 74 *×* 14 = 30,044 combinations. Decomposition reproduces all labeled lipids, and full-space RDKit round-trip validation succeeds. The remaining 28,844 molecules require oracle extrapolation. At *B* = 80, only about 0.27% of this space can be queried. Preliminary experiments also use the original library to assess saturation. A broader data plan includes roughly 1,080 Ugi lipids, within-source rank normalization, source indicators for assay harmonization, and checksum logging.

Random, scaffold-cold, and chemotype-cold splits probe increasing extrapolation. Each uses 960 training and 240 test molecules, the split-generation seed is zero. A held-out Spearman correlation *ρ ≥* 0.3 is the validity gate for each readout. The primary LLM protocol uses eight paired seeds per split, *B* = 40, *m* = 10, *q* = 5, temperature 0.7, and 80 proposal calls. Static-GA, random search, and a non-evolutionary Thompson bandit share the vocabulary, oracle, acquisition, and budget. The bandit refits per-fragment Beta posteriors from the Pareto archive each round. Ablations compare proposal and acquisition alone, Verifier addition, full LEAF, disabled Ledger feedback, and replacement of debate with a binary filter. LipoAgent-related evaluation is restricted to a separate prediction task.

The primary statistic is median paired hypervolume difference, with a 95% percentile bootstrap confidence interval from 10,000 resamples and random seed zero. Two-sided Wilcoxon tests are secondary. Minimum exact *p*-values are 0.25, 0.125, and 0.0078125 for *n* = 3, 4, and 8, eight seeds permit significance but do not guarantee adequate power. The primary multiplicity family comprises four models against two baselines on the random split, with Holm correction. All other comparisons are secondary or exploratory, their displayed *p*-values are unadjusted. Post hoc equivalence tests use a margin of 10% of the baseline median. Full-space normalized hypervolume is defined in Appendix G. Hypotheses H1 predicts higher hypervolume than both baselines, especially on chemotype-cold splits. H2 predicts improved sample efficiency from Ledger feedback, H3 predicts lower false acceptance at matched coverage from recalibrated debate, H4 predicts leakage-conditional rediscovery of the brominated-tail association. An H1 null was internally pre-committed to mean no measured design-efficiency benefit.

## Algorithm 1 Budget-accounted LEAF round

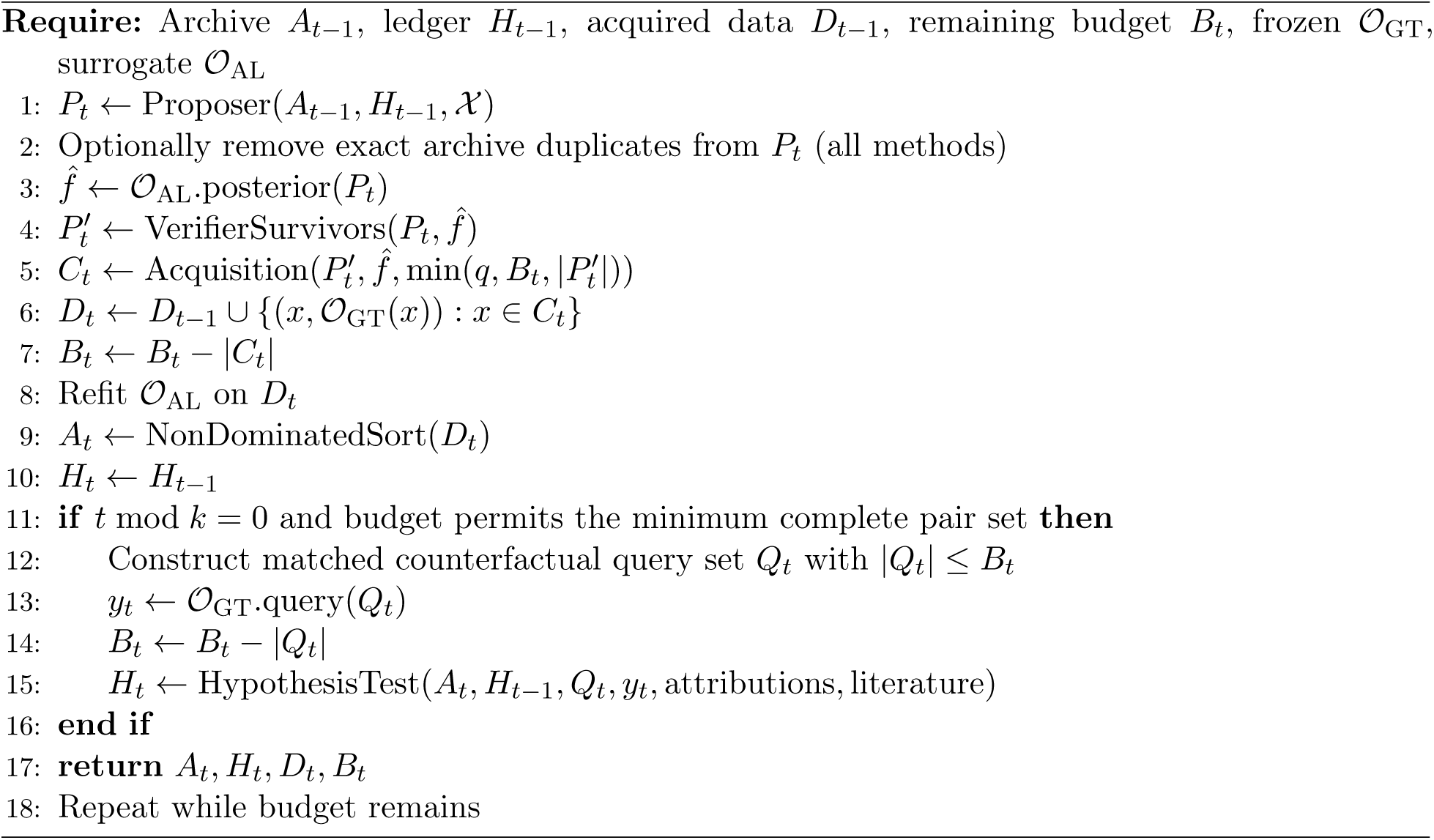

## 3 Results and mechanism analysis

### 3.1 Oracle validity and zero-shot prediction

Held-out HeLa correlations are 0.68, 0.66, and 0.49 on random, scaffold-cold, and chemotype-cold splits, respectively, all above the validity gate. RAW264.7 is weaker and fails on chemotype-cold with *ρ* = 0.23, AGILE contains no toxicity assay. Figure 2A supports controlled oracle comparisons, not biological accuracy for every extrapolated molecule.

#### Validated chemotype-cold endpoint

Repeating the primary Llama 8B protocol with HeLa alone gives median differences of *−*3.93 against both baselines, each with *p* = 0.015625. Baseline hypervolumes are identical across all eight paired seeds under this metric. The two-objective differences of *−*31.2 and *−*46.3 remain diagnostic, but the single validated assay provides the interpretable endpoint. Its roughly 28% deficit relative to a reference hypervolume near 14 preserves the direction at a smaller numerical scale, absolute one- and two-objective hypervolumes are not directly comparable. Appendix J retains per-seed values.

A separate control asks Qwen3-8B, a base model used by LipoAgent (Li et al., 2026), to score efficiency from 1 to 10 at temperature zero without labels. On 50 held-out random-split lipids, rejection is zero and potency correlation is negligible: *ρ* = 0.014, *p* = 0.92, versus the reported fitted-oracle correlation of 0.68 on the matched molecules. This does not replicate LipoAgent’s fine-tuning, toxicity conditioning, prompts, TxGemma component, or ensemble and cannot refute the complete system.

### 3.2 Matched-budget proposal comparisons

An archive-guided rule-based stand-in is not an LLM. At *B* = 400 on 1,200 candidates, random search nearly exhausts the frontier, with zero chemotype-cold hypervolume standard deviation and all split-wise *p ≥* 0.17. After correcting two implementation errors, reducing *B* to 80 yields stand-in differences from GA of 5.5, *−*11.9, and 0.4, with *p* = 0.95, 0.20, and 0.74. Expanding to 30,044 candidates also gives no significant gain. The *p*-values are 1.0, 0.84, and 0.87 against GA and 0.74, 0.46, and 0.74 against random search. Thus saturation alone does not explain the negative results.

Table 1 preserves all split-wise medians and tests. Llama 8B and Qwen 7B have negative differences in all 12 comparisons, with four and three unadjusted significant tests, respectively. Qwen 14B, evaluated on a 16 GB RTX A4000, shows no detectable difference from either baseline on random or scaffold-cold splits, but equivalence is also unestablished. Its chemotype-cold differences are *−*55.1 and *−*77.0, both *p* = 0.0078, with hypervolume confined to 84.7-84.9 across eight seeds. Flat oracle responses are a plausible, unverified explanation in this weak-validity regime. Gemini Flash-Lite has negative differences against both baselines on all splits, each *p* = 0.0078, with zero rejection.

**Table 1:** Eight-seed proposal comparisons. Each cell reports the median paired hypervolume difference from baseline, with the unadjusted two-sided Wilcoxon *p*-value in parentheses. GA denotes static-GA, RS denotes random search. Chemotype-cold two-objective results are diagnostic because one assay fails validation. Exact diversity-prompt tests use the same per-seed results as the other comparisons.

| Method | Baseline | Random | Scaffold-cold | Chemotype-cold |
| --- | --- | --- | --- | --- |
| Llama 3.1 8B | GA | −47.1 (0.055) | −24.8 (0.078) | −31.2 (0.039) |
|  | RS | −42.0 (0.0078) | −28.8 (0.039) | −46.3 (0.016) |
| Qwen2.5 7B | GA | −40.9 (0.078) | −21.6 (0.109) | −29.2 (0.016) |
|  | RS | −29.1 (0.016) | −34.3 (0.195) | −43.4 (0.023) |
| Qwen2.5 14B | GA | +1.8 (0.742) | +18.3 (0.313) | −55.1 (0.0078) |
|  | RS | −4.3 (0.742) | +33.9 (0.250) | −77.0 (0.0078) |
| Gemini Flash-Lite | GA | −91.8 (0.0078) | −66.6 (0.0078) | −24.8 (0.0078) |
|  | RS | −85.5 (0.0078) | −71.1 (0.0078) | −60.2 (0.0078) |
| Diversity prompt | GA | −32.5 (0.078) | −15.6 (0.148) | −10.1 (0.25) |
|  | RS | −25.4 (0.016) | −18.0 (0.0078) | −31.5 (0.078) |
| Thompson bandit | GA | −16.2 (0.742) | +5.9 (0.945) | −18.0 (0.028) |
|  | RS | +0.9 (0.844) | +3.4 (1.000) | −35.2 (0.039) |

Across the 32 random-split model-seed pairs, 24 differences from GA are negative, with median *−*38.7, 27 differences from random search are negative, with median *−*32.6. These are descriptive

The Thompson bandit shows no detectable difference on random or scaffold-cold splits but is worse on chemotype-cold, indicating that cold-split difficulty is not exclusive to LLM reasoning. Groq-hosted Llama 3.3 70B has only three random-split seeds: hypervolumes 107.1, 116.2, and 116.2, versus GA’s 182.1, 112.5, and 135.5 and random search’s 139.1, 155.1, and 166.7. Median differences are *−*19.3, CI [*−*75.0, 3.7], and *−*38.8, CI [*−*50.5*, −*32.0]. These results are exploratory. Rejection is zero, duplication 7.1%, and mean edit distance 1.76. Quota-driven vocabulary subsampling also differs from the primary prompts, limiting scale attribution.

### 3.3 Duplicate queries, novelty, and interventions

#### Proposal pools and selected queries

Initial four-seed diagnostics show archive duplication of 67.8% for Llama 8B and 62.5% for Qwen 7B, with edit distances 0.587 and 0.650, GA gives 22.5% and 1.516, and random search 0% and 2.812. Llama and Qwen label 99.7% and 56.2% of proposals as crossover. Baseline labels differ, so their zero crossover-label rate does not imply no recombination. One eight-candidate Llama pool repeats a triple six times. Direct instrumentation of Llama’s 320 charged random-split queries across eight seeds finds 79.1% exact archive duplicates, versus 15.3% for GA and 0% for random search. Acquisition therefore does not eliminate repetition. Known-good duplicates may inherit attractive surrogate means, although this mechanism is not separately established. Querying the first *q* verified candidates without UCB gives no measurable improvement, implicating poor pool diversity as well as possible selection effects.

#### Applicability domain

Among those Llama queries, 10.3% contain no reagent unseen in oracle training, 88.75% contain one, 0.9% contain two, and none contain three. Mean unseen-slot counts are 0.91 for Llama, 1.58 for GA, and 1.81 for random search. Archive-anchored proposals therefore extrapolate less than full-vocabulary baselines, whose draws encounter many added aldehydes. Greater measured extrapolation does not explain Llama’s deficit in this condition, these counts do not independently establish the cause. This accounting covers one model and split only.

Qwen 14B has 4.2% duplicates on random and scaffold-cold splits and 2.1% on chemotype-cold, with edit distances 1.63 and 1.69. Gemini’s split-wise rates are 43.1%, 34.5%, and 19.2%, with distances 0.90, 0.98, and 1.14. Across 60 diagnosed runs, duplication and deficit are not significantly correlated, *ρ* = *−*0.117, *p* = 0.37. Cross-model averages therefore do not establish a seed-level dose response, novel-candidate quality, acquisition, and oracle extrapolation remain relevant.

#### Prompt and temperature controls

Listing triples to avoid reduces duplication from 67.8% to 34.0% and increases edit distance from 0.59 to 1.06. Across eight seeds on each split, all six median differences remain negative, only the random and scaffold-cold comparisons against random search are significant before multiplicity correction. An exploratory four-seed temperature check gives 87.5% duplication and identical hypervolume 106.4 at temperature 0.2. At temperature 1.0, duplication is 72.2% and median hypervolume 126.3, versus 110.4 for matched default-temperature seeds. This does not establish monotonic improvement, duplication remains above the separate default diagnostic rate.

#### Archive de-duplication

Removing exact current-archive matches before scoring, uniformly across methods and without backfill, improves a four-seed *B* = 15 pilot from *−*17.6 to *−*4.7 against GA and from *−*15.1 to +2.3 against random search, all CIs cross zero. The *B* = 40 replication and cold-split extensions are retained in Table 2. Random-split medians narrow without changing sign, and both unadjusted tests become non-significant. Its independent control differs from the primary run in magnitude, consistent with stochastic proposal variation, the GA comparison also crosses the significance threshold. Scaffold-cold narrows substantially, whereas chemotype-cold changes less, so duplication is not a uniform explanation. The eight-seed de-duplicated arm requires about ten hours. Seeds 4 and 7 stop after only 31 and 14 queries when repeated empty pools trigger the safety limit, all cold-split seeds spend *B* = 40. Equal budget caps therefore do not imply equal realized query counts. De-duplication increases proposal cost and can prevent budget exhaustion, the pilot already required 2.3-7 times as many rounds.

**Table 2:** Matched archive de-duplication at *B* = 40. Entries give median paired hypervolume differences against GA and random search, in that order. Random-split comparisons use *n* = 8 seeds. Scaffold-cold and chemotype-cold are exploratory extensions at *n* = 4. Two random-split de-duplication seeds do not exhaust the budget.

| Split | $n$ | Control: GA, RS | De-duplicated: GA, RS |
| --- | --- | --- | --- |
| Random | 8 | -33.0, -38.7 | -21.9, -35.7 |
| Scaffold-cold | 4 | -34.2, -40.0 | -5.1, -13.4 |
| Chemotype-cold | 4 | -45.3, -75.1 | -35.0, -70.2 |

#### All-evaluated de-duplication

Archive-only filtering permits repeat queries to dominated candidates and duplicate novel proposals within a pool. A stricter control removes both, again without backfill, subsequent rounds replenish the pool. On random-split seeds 0-3, it reverses the median against GA and reduces the random-search deficit by roughly 70% (Table 3). CIs remain wide and cross zero: [*−*46.6, 119.8] against GA and [*−*32.2, 77.2] against random search. This is directional evidence that novelty control matters, not superiority at *n* = 4.

**Table 3:** All-evaluated de-duplication on four matched random-split seeds. Values are median paired hypervolume differences.

| Baseline | No de-duplication | All-evaluated de-duplication |
| --- | --- | --- |
| Static-GA | −33.5 | +20.1 |
| Random search | −52.5 | −15.4 |

#### Objective-scored prompt

The default prompt exposes fragment identities but neither potency values nor an explicit objective description. Adding both assay scores for sampled archive members and an instruction to maximize both objectives yields Table 4. The GA median becomes positive and the random-search deficit narrows by about 38%, CIs are [*−*63.2, 119.8] and [*−*42.9, 77.2], respectively. Seed 1 reaches hypervolume 232.263 in both this experiment and stricter de-duplication, agreeing to twelve significant figures despite distinct query trajectories. This rules out identical cached trajectories but makes the checks dependent through a shared starting seed and convergent endpoint. The two exploratory improvements are not independent replications. Combining scored prompts with novelty control and repeating at *n ≥* 8 remain future work.

**Table 4:** Objective-scored prompting on four matched random-split seeds. Values are median paired hypervolume differences, both intervention intervals cross zero.

| Baseline | Default prompt | Objective-scored prompt |
| --- | --- | --- |
| Static-GA | −33.5 | +13.9 |
| Random search | −52.5 | −32.7 |

#### Practical cost

Candidate latencies are approximately 4.0, 10.3, 10.3, and 5.1 seconds for Llama 8B, Qwen 7B, Qwen 14B, and Gemini. Groq takes about 0.6 seconds of raw inference but is paced at seven seconds. Non-LLM methods require no LLM calls, the recorded non-LLM runs and GP rounds each take under one second. These workload-specific timings exclude quotas, retries, and multiday accumulation beyond nominal 80-call costs.

### 3.4 Does the proposer genuinely use injected evidence?

The objective-scored prompt (Table 4) could improve either because the model exploits the injected potency values or because it already carries exploitable pretrained knowledge, or because any visible signal, true or not, shifts behavior without being used as intended. The following six checks triangulate this question. Appendix K–K.6 give full per-seed data.

#### Leakage probe

Llama 3.1 8B, asked to zero-shot score the same 50 held-out random-split lipids as the Qwen3-8B control (Section 3.1) without labels, returns the identical score (8/10) for all 50 molecules at temperature zero, a degenerate constant output for which Spearman correlation is undefined. Repeating at temperature 0.7 breaks the degeneracy (three distinct emitted scores, 6-8) and gives *ρ* = *−*0.006, *p* = 0.96, confirming the null is a genuine absence of exploitable pretrained signal rather than a greedy-decoding artifact, and consistent with Qwen3-8B’s own near-zero *ρ* = 0.014 control.

#### Rank-only prompt

Replacing explicit potency and RAW264.7 magnitudes with ordinal rank 1-8 among the shown archive members, removing numeric values entirely, yields medians of +17.2 against GA and *−*17.7 against random search on four matched random-split seeds. This represents a larger GA-relative gain and a smaller random-search-relative deficit than the full-magnitude scored prompt (Table 4). Ordinal information alone is therefore sufficient to recover, and on this metric exceed, the scored-prompt effect, which does not require exact numeric values. Both CIs cross zero at *n* = 4.

#### Chain-of-thought rationale

Requiring a one-to-two sentence structural rationale before each proposal, enforced by rejecting outputs lacking one, gives medians of *−*6.4 against GA and *−*43.3 against random search, within the range of the other prompt variants and not a qualitative improvement over the default prompt.

#### Score-shuffle perturbation

Following Fang et al. (2026), real potency values are retained but randomly reassigned to the wrong fragments before display. Aggregate median differences (+13.9 GA, *−*32.7 RS) are essentially identical to the correctly labeled scored prompt (Table 4: +13.9, *−*32.7). Per-query comparison against the matched correctly scored run shows this is not because the proposer ignores the (now-false) scores: 28, 8, 0, and 7 of 40 queries differ by seed, so three of four seeds have substantial real divergence while one seed’s trajectory is fully identical. The narrow attainable Pareto frontier lets different query trajectories, including ones built on false score mappings, converge on the same endpoint hypervolume, so aggregate equivalence under shuffled scores does not by itself demonstrate that scores are ignored.

#### Evidence-conflict probe

A fixed eight-member archive sample is shown with deliberately inverted rank labels (true best relabeled worst and vice versa) in a rank-only prompt, and 40 independent Llama 8B calls are scored for which archive member the proposal is nearest to by fragment edit distance. Repeating this with three independently drawn archive samples reveals the result is highly sample-dependent rather than a single stable behavior. Resolved calls following the falsely labeled-best member (the true worst) are 14.8% (4/27 resolved), 84.2% (32/38), and 46.2% (6/13) across the three samples, with correspondingly variable tie/unresolved rates (13, 2, and 27 of 40 calls). One pattern is consistent across all three samples. The genuinely best member is almost never the nearest neighbor regardless of which label it carries, with 0, 3, and 0 resolved calls across the samples. The pooled rate is 3/78, or 3.8%, versus a 12.5% uniform base rate. Beyond that one robust null, a single fixed-archive run cannot characterize whether or how strongly the proposer follows a falsified quality label. The earlier characterization of this probe as showing “list-position dominance” was based on one sample and does not generalize. The only well-supported claim is that true quality is not what drives these proposals, not a specific alternative mechanism.

#### Genuine reasoning-mode proposer

Enabling Qwen3-8B’s native extended-thinking mode (not the prompted v5_cot rationale) produces real multi-thousand-character reasoning traces under the default (unscored) prompt (mean length 5,669 characters, present in 155/160 calls) that never mention potency or the objective (0/155). Under the scored prompt, explicit engagement rises sharply to 124/160 traces (77.5%) mentioning potency, with 152/160 containing a trace (mean length 6,877 characters). Median hypervolume differences are *−*19.1 against GA and *−*53.4 against random search on four seeds. These results are weaker than those for non-reasoning Llama 8B under the identical scored prompt (Table 4), and the proposer is 5-=-10*×* slower per call. Genuine engagement with injected evidence is therefore dissociable from search performance.

### 3.5 Structural inspection: best-found lipids and fragment associations

The preceding sections report only aggregate hypervolume. Figure 5A shows what the search loop actually finds. For the objective-scored-prompt check’s seed 1 (Section 3.3), we identify each method’s single best-found candidate among its 40 queried triples, scored by the same frozen *O*_GT_ used throughout, and render its real assembled structure. All three methods converge on branched tertiary-amine head groups and long aliphatic tails, the qualitative lipid-like shape AGILE’s chemistry space is built to produce (Xu et al., 2024). Llama 8B’s best candidate has the highest predicted sum (19.3 vs. 15.2 for random search and 11.5 for GA) on this one seed, consistent with that seed’s favorable scored-prompt hypervolume (Table 4) but not a general claim about structure quality across seeds.

A separate, fully post-hoc diagnostic is explicitly independent of the Hypothesis Ledger, which this paper documents as empty in every search run where it was enabled (Section 3.6). The diagnostic inspects what the frozen random-split *O*_GT_ associates with higher predicted potency across the complete 30,044-candidate space, with no search, no budget charge, and no counterfactual-pair construction. Figure 5B shows the five highest- and lowest-scoring values at each reagent slot by mean predicted HeLa+RAW264.7 across every candidate containing that value. The isocyanide slot (C) shows a narrow range (6.83-7.38) dominated by tail length, with the two mono-unsaturated tails scoring marginally highest. The aldehyde slot (B) shows the opposite relationship. Fully saturated, branched linkers score highest, and doubly unsaturated (diene) tails score lowest. This directly contradicts a simple “unsaturation improves potency” reading if that were inferred from slot C alone. The inconsistency across reaction positions is the main finding. It indicates that the oracle’s fitted structure-potency relationship is specific to each slot’s chemical context rather than a single interpretable rule. It also explains why this result is reported as a retrospective correlational inspection of the fitted surrogate rather than a validated structure-activity hypothesis or a replication of the Ledger’s intended matched-pair statistical test. No halogenated reagents exist in this vocabulary, so this analysis cannot speak to the LUMI-lab brominated-tail association (Xu et al., 2026) discussed in Appendix A.

#### Domain-coverage boundary

None of the clinically deployed ionizable lipids examined here— MC3 (DLin-MC3-DMA), SM-102, and ALC-0315 (Escalona-Rayo et al., 2023)—decompose into this paper’s Ugi-3CR vocabulary under the same retro-synthetic decomposition routine validated to exactly round-trip all 1,200 labeled AGILE lipids (Section 2.3, Appendix L). This is expected because none are Ugi three-component-reaction products. It converts the planned but previously unreported Recall@*k* comparison against these reference compounds (Section 2.3) into an explicit, disclosed boundary of this chemistry’s coverage rather than a silently absent result. The search space studied here is disjoint from these clinically deployed lipid chemotypes. No claim in this paper about proposal quality should be read as bearing on recovery of, or comparison to, these specific compounds.

### 3.6 Secondary-operator ablations and implementation audit

Ledger and binary-filter ablations show no significant hypervolume change. At *B* = 80, *m* = 40, *q* = 8, and *n* = 8, disabling the Ledger on chemotype-cold or replacing verification with a binary filter on the random split yields results identical to full LEAF. The Ledger is empty in all 72 original sweep runs, comprising three methods, three splits, and eight seeds, hypothesis precision is undefined, not zero. A reproduced *B* = 40, *q* = 5, *k* = 4 run has 12 Pareto members sharing an isocyanide different from the fixed pairing control. Archive-restricted matching prevents completed tests despite passing validation unit tests. Deterministic toxicity likewise makes debate and binary filtering equivalent, with undefined binary-filter coverage. These outcomes reflect the absence of usable signal under the tested conditions, not an evaluation of effective debate or hypothesis feedback.

An early stand-in result gave a median GA advantage of 92.5, CI [30.1, 119.1], *p* = 0.0078. Reusing a full-data oracle invalidated split distinctions, while GA’s mutation probability 0.2 was unmatched to the stand-in’s 50% pure mutation. Correcting both erased the effect. A third error prevented EHVI dispatch, historical runs actually used UCB with the intended *β* = 2. After correction, exploratory EHVI on four random-split seeds gives differences of *−*23.5 against GA and *−*38.2 against random search, similar in direction and magnitude to UCB. The exact four-seed test cannot attain *p <* 0.05, an all-split *n ≥* 8 EHVI sweep remains necessary.

## 4 Discussion, limitations, and conclusion

No tested configuration establishes an advantage for LLM proposers under matched query budgets. Small models waste a large fraction of evaluations on exact duplicates. Larger models reduce this repetition yet still fail to outperform genetic-algorithm or random-search baselines. Prompt and novelty interventions can shrink the gap, but only at higher proposal cost, so efficiency and latency must be judged together. Statistically, none of the eight primary comparisons survives Holm correction, and post-hoc equivalence tests at a 10 % margin also fail. Non-significance therefore shows neither inferiority nor parity. Small samples, shared baselines, and model heterogeneity further limit formal claims.

The study is model-on-model: search trajectories are scored by a frozen Gaussian-process oracle rather than wet-lab measurements, so results depend on the fidelity of that surrogate and on the particular lipid space examined. Chemotype-cold validity is weak, toxicity screening is purely rule-based, and the expanded 30,044-candidate library requires extrapolation. The Verifier is active in every reported configuration but is always degenerate in this dataset, since AGILE supplies no independent toxicity assay. Its conformal intervals have zero width, and its decisions reduce to a deterministic rule. It has not been exercised under genuine epistemic uncertainty. The Hypothesis Ledger is active in the primary sweep and early ablations, where it is measured empty across all 72 runs because the Pareto-restricted counterfactual-pair construction rarely finds a matched control. It is explicitly disabled in essentially every later follow-up experiment, including every check in Section 3.4, rather than silently inactive. Coverage is limited to four models, three splits, and a modest set of prompt and acquisition variants. Several follow-up checks remain under-powered, and access constraints on larger or domain-adapted models restrict broader comparison.

Under controlled oracle access, identical acquisition functions, and matched budgets, the tested language-model proposers do not improve lipid search relative to classical baselines. The dominant inefficiency is duplicate-driven query waste, which lessens with scale but is not replaced by a performance gain. LEAF’s contribution is a reusable matched-budget evaluation protocol and a concrete diagnosis of where current zero-shot LLM proposers lose efficiency, not a claim about biological performance or the ultimate limits of LLM-driven methods more broadly.

## Data availability

The labeled AGILE dataset and its unlabeled virtual candidate library originate from the AGILE platform’s own repository at https://github.com/bowang-lab/AGILE (Xu et al., 2024).

## Code availability

All code used to run experiments, compute statistics, and generate figures is available at https://github.com/harshitsinghsnu/LEAF.

## Acknowledgments

Harshit Singh acknowledges the support of the Department of Computer Science and Engineering, Shiv Nadar University, Delhi-NCR, for providing the computational resources used in this work. Rohan Gorantla acknowledges the support and funding of United Kingdom Research and Innovation (grant EP/S02431X/1), the UKRI Centre for Doctoral Training in Biomedical AI at the School of Informatics, University of Edinburgh, and Exscientia Plc, Oxford.

## A Secondary-agent specification and implementation scope

### A.1 Debate and conformal verification

The general Verifier comprises *N* independently prompted toxicologists spanning at least two base models. Each uses RDKit toxicophore matching, an active-learning toxicity posterior when available, and free-text reasoning. Same-model prompt or temperature variants measure prompt sensitivity rather than independent epistemic diversity. Inter-agent agreement above 95% flags insufficient diversity for H3.

Held-out split-conformal calibration converts aggregated scores to intervals (Angelopoulos and Bates, 2023), intervals crossing the decision threshold trigger abstention. Adaptive acquisition and refitting invalidate ordinary exchangeability. Weighted or Mondrian recalibration therefore occurs each round, with empirical coverage monitoring rather than a distribution-free guarantee under arbitrary shift (Tibshirani et al., 2019; Vovk, 2013). Calibration provenance, weighting, strata, and thresholds must be specified.

The AGILE experiments contain no learned toxicity posterior. Calibration scores equal deterministic toxicophore labels, so residual uncertainty and interval width are zero. Repeated rule outputs reproduce binary-filter decisions, not independent epistemic evidence or demonstrated calibration of a learned safety signal. Testing this component requires independent toxicity labels and assay harmonization.

### A.2 Hypothesis generation and testing

Every *k* rounds, indexed LNP literature and Integrated Gradients or KernelSHAP attributions over the Pareto archive motivate structure-property hypotheses. Each rule requires matched counterfactual pairs, a minimum pair count, and a paired Wilcoxon test. Only frozen *O*_GT_ supplies validation labels. All queried molecules, including controls, consume the shared budget. Incomplete pair sets remain untested, the Ledger updates only after validation and then informs proposals when enabled. Algorithm 1 bounds complete pair sets by the remaining budget.

Pair construction searches only Pareto members and requires a fixed control reagent. In the reproduced *B* = 40, *q* = 5, *k* = 4 configuration, all 12 final members share one isocyanide among 14 possible values, different from that control. Correlated objectives can concentrate the frontier and eliminate matched controls. This explains the empty Ledger and undefined precision across 72 original sweep runs. Passing validation-logic unit tests does not establish that the optimization loop supplies testable pairs.

The LUMI-lab association concerns brominated tails and potency, not organ tropism (Xu et al., 2026). Because its preprint was public from February 2025, a bare-LLM probe without retrieval or archive context must precede any rediscovery claim, with either outcome disclosed. Detected prior knowledge changes the interpretation to consistency checking. Neither the original AGILE vocabulary nor its virtual-library extension supplies halogenated reagents. H4 therefore requires literature-sourced, Ugi-compatible halogenated reagent structures and was not run.

## B Hosted inference and resource accounting

Groq uses a chat-completions interface and Gemini a native interface. Both obtain credentials from environment variables, not stored experiment settings or command-line arguments. The following measurements describe the experimental period, not current quotas, prices, or availability.

**Groq.** The recorded allowance for llama-3.3-70b-versatile was 100,000 tokens per day. Full-vocabulary prompts exposed 29 amines, 74 aldehydes, and 14 isocyanides and consumed about 1,850 tokens per call, or 148,000 per 80-call seed before retries. A second independent key encountered the same ceiling. Sampling eight values per slot reduced consumption to about 813 tokens per call and 65,000 per seed. Output validation still used the complete vocabulary, preserving validity but changing each call’s visible choices.

The allowance recovered over an observed rolling 24-hour window. Three daily attempts failed before a fourth succeeded, large quota probes were replaced by trivial probes to conserve tokens. Three random-split seeds accumulated over several days, all with zero rejection. Two underperformed both baselines, one exceeded GA only. A later catalog query reportedly omitted the model, but no dated endpoint snapshot supports a present-availability claim. An eight-seed, three-split sweep was not completed.

**Gemini.** The recorded gemini-flash-latest allowance was 20 requests per day in its free-tier per-project, per-model bucket. The separate gemini-flash-lite-latest bucket enabled eight seeds on all three splits. These completed results supersede partial runs. Flash-Lite is not the flagship model or an established frontier-capability benchmark.

**Rate limiting.** Hosted calls are paced about seven seconds apart. HTTP 429 responses invoke Groq’s Retry-After header or an empirical Gemini delay, with a configured retry limit of eight before raising an error rather than silently replacing the model. Pacing controls bursts, not daily token totals, compression and quota recovery address the latter. Nominal workload costs exclude multiday scheduling and some retries.

**Domain-adapted models.** TxGemma was not evaluated because of access gating and insufficient 16 GB VRAM for its 27B chat model. Prebuilt ChemLLM and ChemDFM checkpoints were unavailable in the local backend. Meditron 7B, continued-pretrained on biomedical and clinical text from Llama 2, repeatedly returned unrelated tool-shaped JSON, invalid reagent structures, or literal vocabulary placeholders. Its 100% rejection and fallback rate precluded a meaningful sweep. Per-run rejection logging distinguishes this failure from genuine model performance.

## C Diagnostic example and experiment provenance

### C.1 Repeated proposals within one pool

A default-prompt Llama 8B pool of eight candidates, initialized from the recorded six-fragment random archive, repeats the first triple six times and each remaining triple once. All are labeled crossover. Ellipses in the last two isocyanide slots are retained abbreviations, not reconstructed molecules.

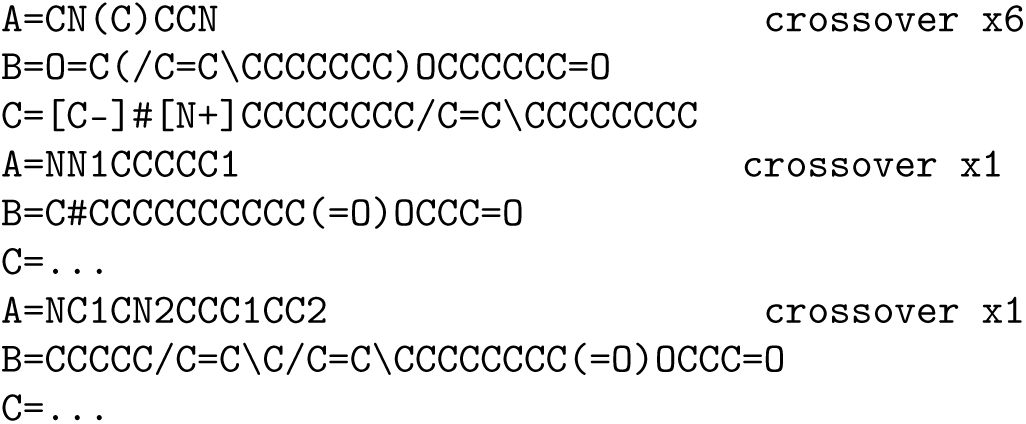

Within-pool repetition differs from archive duplication. Archive-only de-duplication removes existing Pareto matches without backfill but does not ensure mutual uniqueness among novel proposals.

### C.2 Scope and reproducibility

The experiment record distinguishes oracle-validation outputs, per-seed search results, acquisition-disabled diagnostics, diversity prompting, and the corrected acquisition dispatch. These provenance records describe the study, the manuscript package does not contain an anonymous code or raw-trace release. Initial Llama and proposal-diversity diagnostics used four seeds before the eight-seed, three-split sweep. Later panel aggregates summarize larger samples and need not equal pilot rates. Acquisition-disabled results are diagnostic, not a complete eight-seed factorial study.

Budget regimes are sequential controls, not interchangeable descriptions: *B* = 400 tests saturation on the original 1,200-candidate space, *B* = 80 reduces the budget before and after expansion to 30,044 candidates, *B* = 40 defines the primary LLM protocol, and *B* = 15 defines the initial four-seed de-duplication pilot. Primary inference uses eight random-split comparisons. Other tests are secondary or exploratory, including eight-seed de-duplication.

The recorded reproducibility materials include exact partition indices with split-generation seed zero and 960 training and 240 test molecules per split, plus a manifest mapping tables to experiment settings and outputs. GP settings are a Tanimoto kernel, diagonal regularization *α* = 10*^−^*^6^, internal target normalization, and random seed zero. Bootstrap intervals use 10,000 resamples, 95% two-sided percentile bounds, and random seed zero. Objective definitions and the single-assay exception appear in Section 2.1. The internal null-result interpretation was not independently timestamped, a complete anonymous code and trace release remains separate from this manuscript.

### C.3 Eight-seed de-duplication and exploratory EHVI

For the random-split *B* = 40 archive-de-duplication intervention, Llama 8B hypervolumes are 135.463, 90.001, 134.490, 123.479, 100.779, 100.779, 167.939, and 100.779. Matched GA values are 159.384, 109.838, 135.463, 114.082, 131.714, 201.687, 134.490, and 152.093, random-search values are 139.085, 155.060, 166.663, 162.617, 124.540, 175.418, 130.415, and 155.060.

The independently replicated no-de-duplication control gives Llama values 100.779, 80.137, 100.779, 123.479, 100.779, 100.779, 100.779, and 128.397. Its GA values are 159.384, 112.501, 135.463, 100.518, 125.852, 199.167, 134.490, and 144.025, the random-search vector is unchanged. De-duplication applies to every method, so GA’s nonzero duplication changes its trajectory, whereas no observed random-search duplicates are removed. Using the distinct GA control reproduces the median *−*33.0 and *p* = 0.023 in Table 2. A prior transcription incorrectly reused the de-duplicated GA vector for this control, the corrected vectors retain the reported summary statistics.

De-duplicated seeds 4 and 7 terminate after 31 and 14 queries because of repeated empty pools, not runtime errors. Thus the nominal *B* = 40 cap is not exhausted in these two cases.

The four-seed EHVI run gives Llama values 100.779, 88.219, 100.779, and 123.479, GA values 123.479, 112.501, 135.463, and 112.282, and random-search values 139.347, 155.060, 132.080, and 161.350. No Wilcoxon result is reported: the recorded invocation failed on the observed tie pattern, and an exact two-sided four-seed test cannot attain *p <* 0.05 even without ties. Small *n* limits significance resolution, rather than making every signed-rank calculation intrinsically invalid.

### C.4 Cold-split de-duplication and secondary ablations

At *n* = 4, scaffold-cold control differences are *−*34.22 against GA and *−*39.96 against random search, de-duplication gives *−*5.09 and *−*13.37. Chemotype-cold controls give *−*45.29 and *−*75.06, changing to *−*34.98 and *−*70.16. All seeds exhaust *B* = 40. No signed-rank results are reported for these exploratory comparisons, only median differences and bootstrap intervals are used. Scaffold-cold narrows by factors of about 6 and 3, compared with 1.3 and 1.07 for chemotype-cold. This asymmetry remains mechanistically unresolved and must be interpreted alongside chemotype-cold’s failed RAW264.7 validity gate.

The *B* = 80, *m* = 40, *q* = 8 Ledger and binary-filter ablations cover chemotype-cold and random splits, respectively, at eight seeds, not a complete cross-product. Recorded seed-zero traces report eight Ledger-ablation rounds with six final-round queries and nine binary-filter rounds with four final-round queries. These counts alone cannot reconcile *B* = 80 with at most eight selected queries per round: partial final rounds do not explain the discrepancy. Complete initialization and query accounting are required before these trace counts can substantiate budget exhaustion. This limitation does not alter the reported equality of ablation outcomes.

### C.5 Reporting corrections

Llama has four unadjusted significant comparisons, not five, together with Qwen 7B this is seven of twelve. Qwen 14B’s positive random-split point is against GA. Groq’s 7.1% duplication is below the small-model and GA rates but above Qwen 14B’s aggregate 3.5%. Temperature 1.0’s 72.2% duplication is not below the separate 67.8% default diagnostic. Seven to 21 de-duplicated pilot rounds versus three control rounds imply a 2.3-7-fold increase. The three implementation errors concern split-oracle reuse, mutation mismatch, and acquisition dispatch. These distinctions correct interpretation without changing measured entries.

## D Baseline proposal-pool specification

All methods share archive and vocabulary access, *m* = 10, *q* = 5, the evaluation oracle, and the acquisition function. By default, no method removes within-pool duplicates. Archive-only filtering, when enabled, removes exact Pareto matches before scoring, all-evaluated filtering also removes previously evaluated dominated candidates and within-pool repetitions.

Each method constructs its proposal pool differently from a common archive and vocabulary. The LLM proposer draws up to eight sampled Pareto members per call, mutates by having the model choose one slot to change, and crosses over by having the model recombine two archive parents, duplicate handling is none by default, with the common optional filters available to every method. Static-GA draws two uniformly sampled archive members without replacement as parents, mutates with probability 0.5 by redrawing one random slot uniformly, and crosses over by copying each slot from a uniformly chosen parent, duplicate handling is likewise none by default with the same optional filters available. Random search draws independent uniform vocabulary samples with no parent source, mutation, or crossover operator, and no duplicate handling is needed since its measured archive-duplicate rate is zero. The Thompson bandit refits per-slot Beta posteriors from the archive each round, draws independently from those posteriors categorically (its analog of mutation), performs no parent recombination (no crossover), and applies no duplicate handling by default.

Invalid LLM outputs are replaced individually by the rule-based proposer and counted in the rejection rate. By contrast, no method backfills candidates removed by verification or novelty filtering to restore a fixed pool size. If fewer than *q* survive, acquisition queries min(*q, B_t_, |P ^′^|*); subsequent rounds replenish proposals. GA falls back to a uniform triple only with fewer than two archive members, including the initial empty archive. All methods use the same UCB or EHVI selection, not method-specific shortcuts.

## E LLM inference and prompt specification

Instruction-tuned identifiers are llama3.1:8b, qwen2.5:7b, and qwen2.5:14b for local Ollama inference, gemini-flash-lite-latest for Gemini, and llama-3.3-70b-versatile for Groq. Local runs use the backend library’s default GGUF quantization, hosted APIs expose no local quantization control. The recorded aliases do not establish immutable checkpoint revisions.

No system prompt is used. Calls are single-turn and stateless, with context reinserted each time. Default temperature is 0.7, exploratory checks use 0.2 and 1.0. Top-*p*, top-*k*, and repetition penalties retain backend defaults. Local calls request JSON formatting and draw a call seed from the proposer’s seeded generator, reproducibility remains backend-version dependent. Hosted calls request JSON in the prompt without server-side schema enforcement and allow 300 output tokens. The default retry limit is two, increased to eight for hosted runs, quota handling is described in Appendix B.

Reagent strings must match vocabulary entries character-for-character. Invalid outputs invoke individual rule-based fallback and are tracked through the rejection rate, which remains at most a few percent except for Meditron’s 100%. Default prompts expose the full vocabulary, Groq alone samples eight values per slot. Each call shows up to eight uniformly sampled archive members. There is no additional truncation or score summary. Consequently, the default proposer sees identities and archive membership, not objective values or an explicit maximization instruction.

The default user-turn template is reproduced below, with archive members and vocabulary lists interpolated at runtime. Line wrapping is typographic only.

You are the Proposer Agent in an evolutionary lipid-design loop. Candidates are Ugi three-component-reaction lipids, each a triple (A, B, C):

A = amine, one of: […]

B = aldehyde, one of: […]

C = isocyanide, one of: […]

Current Pareto archive (sample):

- A=… B=… C=…

[up to 8 lines]

Propose exactly ONE new candidate via crossover (recombining A/B/C from two archive parents) or mutation (changing exactly one of A/B/C from a single archive parent, or a random triple if the archive is empty). The value of A, B, and C in your output MUST be copied EXACTLY (character for character) from the lists above - do not paraphrase or alter the SMILES strings in any way.

Respond with ONLY a single JSON object shaped like:

{”A”: “<ONE of the A values>”, “B”: “<ONE of the B values>”, “C”: “<one of the C *↪→* values>”,

”provenance”: {”operator”: “crossover” or “mutation”, “parents”: [”…”]}}

The diversity variant adds a paragraph describing repetition and listing up to 20 evaluated triples to avoid, without other changes. The scored variant supplies sampled members’ actual HeLa and RAW264.7 values and explicitly instructs maximization of both objectives.

## F Equivalence and multiplicity

A non-significant difference test does not establish equivalence. The post hoc practical margin is 10% of the baseline’s median hypervolume on the relevant split. Table 5 reports the six available two-one-sided-test (TOST) results, it is not a complete equivalence analysis of every primary comparison.

**Table 5:** Post hoc equivalence tests with a margin of 10% of baseline median hypervolume. No reported test reaches *p <* 0.05.

| Model | Baseline | Threshold (±) | Mean diff | TOST $p$ |
| --- | --- | --- | --- | --- |
| Llama 3.1 8B | GA (random) | 15.3 | -45.4 | 0.947 |
| Llama 3.1 8B | RS (random) | 15.5 | -39.2 | 0.988 |
| Qwen2.5 14B | GA (random) | 15.3 | -9.6 | 0.354 |
| Qwen2.5 14B | RS (random) | 15.5 | -3.4 | 0.102 |
| Qwen2.5 14B | GA (scaffold-cold) | 13.7 | +20.6 | 0.630 |
| Qwen2.5 14B | RS (scaffold-cold) | 13.7 | +25.8 | 0.710 |

Qwen 14B therefore has neither evidence of a difference nor evidence of equivalence on random and scaffold-cold splits at this margin and *n* = 8. Larger samples or a different justified margin would be needed to resolve practical equivalence. The margin was not prespecified before experimentation.

The primary multiplicity family contains the four fully replicated models against two baselines on the random split. Table 6 retains the raw *p*-values and adds valid Holm-adjusted values at family-wise *α* = 0.05.

**Table 6:** Holm correction for the eight primary random-split comparisons. GA denotes static-GA and RS random search. The step-down procedure stops at the first failed threshold, tied minimum *p*-values cannot yield an isolated rejection.

| Comparison | Raw $p$ | Ordered threshold | Adjusted $p$ |
| --- | --- | --- | --- |
| Llama 8B vs. RS | 0.0078125 | 0.00625 | 0.0625 |
| Gemini vs. GA | 0.0078125 | 0.00714 | 0.0625 |
| Gemini vs. RS | 0.0078125 | 0.00833 | 0.0625 |
| Qwen 7B vs. RS | 0.015625 | 0.01000 | 0.078125 |
| Llama 8B vs. GA | 0.0546875 | 0.01250 | 0.21875 |
| Qwen 7B vs. GA | 0.078125 | 0.01667 | 0.234375 |
| Qwen 14B vs. GA | 0.7421875 | 0.02500 | 1.0000 |
| Qwen 14B vs. RS | 0.7421875 | 0.05000 | 1.0000 |

**Table 7:** Attainable hypervolume over the complete candidate space under each split-specific frozen oracle.

| Split | Full-space hypervolume |
| --- | --- |
| Random | 258.2 |
| Scaffold-cold | 235.9 |
| Chemotype-cold | 199.5 |

The minimum raw *p* = 0.0078125 exceeds the first threshold, 0.00625, so no null hypothesis is rejected. A previous rank-by-rank interpretation incorrectly retained the third tied comparison while ignoring the step-down stopping rule. Correcting that interpretation weakens model-specific inferiority claims but leaves the conclusion of no demonstrated improvement unchanged. Displayed unadjusted tests elsewhere must not be read as family-wise evidence.

## G Normalized hypervolume

Full-space hypervolume is obtained by embedding and evaluating all 30,044 candidates with each frozen oracle, without LLM calls or a search budget. It is a diagnostic denominator, not information exposed to a budgeted search. With an empty initial archive and initial hypervolume zero,

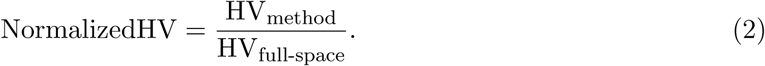

In the primary random-split runs, Llama 8B reaches 106.4-129.5, or 41-50% of the full-space value, GA reaches 103.8-219.7, or 40-85%, random search reaches 124.5-175.4, or 48-68%. These ranges refer to the primary runs, not intervention replicates. They do not establish a universal half-frontier upper bound for all models or interventions. At *B* = 40, the query cap covers about 0.13% of the space. Multiplying every paired difference on a split by the same positive normalization factor changes its scale, not the corresponding significance result.

## H All-evaluated de-duplication

For random-split seeds 0-3 with *B* = 40, stricter de-duplication gives Llama 8B values 135.463, 232.263, 134.490, and 135.463, GA values 182.084, 112.501, 135.463, and 94.317, and random-search values 139.085, 155.060, 166.663, and 162.617. The matched control gives Llama values 100.779, 80.137, 100.779, and 123.479, GA values 159.384, 112.501, 135.463, and 100.518, and the same random-search vector. The unchanged random-search outcomes apply to this matched comparison, not every separate experiment on the split. Seed 1’s value of 232.263 is retained and strongly influences the exploratory result.

## I Objective-scored prompting

The four scored-prompt hypervolumes are 96.167, 232.263, 135.463, and 128.397, at *B* = 40 on the random split. GA values are 159.384, 112.501, 135.463, and 100.518, random-search values are 139.085, 155.060, 166.663, and 162.617. These are the no-de-duplication baseline vectors, not the stricter-filter GA vector. The default-prompt Llama values are 100.779, 80.137, 100.779, and 123.479. Seed 1 reaches 232.262563068695 in both scored prompting and all-evaluated de-duplication. The recorded per-round reagent sequences differ, excluding identical cached query trajectories.

Convergence from a shared seed makes these two exploratory checks dependent, it does not justify treating their favorable medians as independent corroboration.

## J HeLa-only chemotype-cold reanalysis

The single-objective restriction uses the primary *n* = 8, *B* = 40, *m* = 10, *q* = 5 protocol, the HeLa readout, and reference point (*−*5.0). Llama 8B hypervolumes are 10.994, 10.609, 10.527, 10.905, 8.826, 11.422, 12.917, and 11.422. Both baselines give 13.878, 14.147, 17.542, 17.159, 13.142, 16.706, 12.766, and 14.959. Their identical vectors reflect equal best-found values under this metric. Llama’s paired median difference is *−*3.93 against each, with unadjusted *p* = 0.015625. The implementation explicitly makes the objective heads configurable for this exception rather than retaining the historical two-head default.

## K Evidence-use checks

### K.1 Leakage probe (Llama 3.1 8B)

Llama 3.1 8B zero-shot scores the same 50 held-out random-split lipids used for the Qwen3-8B control, from 1 to 10, with no labels. At temperature zero, rejection is zero but all 50 emitted scores equal 8.0 - a degenerate constant output for which Spearman correlation against the fitted oracle’s potency prediction (itself *ρ* = 0.683 on this split, Section 3.1) is undefined rather than computably near zero, this is tracked explicitly rather than silently reported as a correlation of zero. Repeating the identical protocol at temperature 0.7 (same 50 molecules, zero rejection) breaks the degeneracy: three distinct scores are emitted (6, 7, 8), and Spearman correlation is *ρ* = *−*0.0065, *p* = 0.96. This confirms the temperature-zero degeneracy was a greedy-decoding artifact of this specific prompt and model, not evidence that the question cannot be meaningfully answered, and the non-degenerate result is consistent with the separate Qwen3-8B zero-shot control (*ρ* = 0.014, *p* = 0.92, *n* = 50): both models show a measured near-zero correlation once decoding is non-degenerate, supporting the conclusion that no exploitable pretrained potency signal is recoverable from either model under zero-shot scoring.

### K.2 Rank-only prompt (v4_rank)

The prompt lists up to eight sampled archive members in rank order by summed potency and RAW264.7 score, with no numeric magnitudes shown, and otherwise matches the default template. On four matched random-split seeds at *B* = 40, hypervolumes are 128.397, 227.271, 142.046, and 128.397, against the GA baseline vector 159.384, 112.501, 135.463, and 100.518, and random-search vector 139.085, 155.060, 166.663, and 162.617, giving per-seed differences against GA of *−*31.0, +114.8, +6.6, and +27.9 (median +17.2) and against random search of *−*10.7, +72.2, *−*24.6, and *−*34.2 (median *−*17.7), both CIs cross zero at *n* = 4.

### K.3 Chain-of-thought rationale (v5_cot)

The schema requires a non-empty “rationale” field of one to two sentences describing the structural reasoning behind the proposed crossover or mutation, outputs without one are rejected and retried, counted toward the rejection rate. On four matched random-split seeds at *B* = 40, hypervolumes are 128.397, 106.697, 128.397, and 104.358, giving median differences *−*6.4 against GA and *−*43.3 against random search. Rejection remains under 5%, so the requirement is enforced rather than decorative.

### K.4 Score-shuffle perturbation (v6_shuffled)

The prompt displays real potency and RAW264.7 values exactly as in the scored prompt, but the values are independently shuffled across the displayed fragments before rendering, breaking the true value-to-fragment correspondence while preserving the set of displayed numbers. On four matched random-split seeds at *B* = 40, hypervolumes are 96.876, 232.263, 135.463, and 128.397, giving median differences +13.9 against GA and *−*32.7 against random search - matching the correctly scored prompt’s +13.9 and *−*32.7 (Table 4) to one decimal place. Direct per-query comparison of reagent triples against the correctly scored run’s queries.csv finds real divergence in three of four seeds: seed 0 differs on 28/40 queries, seed 1 on 8/40, seed 2 on 0/40 (fully identical trajectory), and seed 3 on 7/40. The attainable Pareto frontier under this budget and split is narrow enough that distinct query trajectories, including trajectories built from deliberately false score mappings, often converge on the same final hypervolume. Aggregate equivalence therefore reflects frontier concentration, not proof that scores go unused.

### K.5 Evidence-conflict probe

A fixed eight-member sample is drawn from the random-split training partition’s evaluated archive, independently for each of three probe-sampling seeds (0, 1, 2). Within each sample, true rank is computed by summed potency and RAW264.7 score, displayed rank labels are the true ranking inverted (true-best shown as rank 8, true-worst shown as rank 1). This falsified ranking is rendered in the v4_rank template and issued to Llama 3.1 8B 40 times independently per sample (temperature 0.7, distinct call seeds), each call producing one proposal via the proposer’s own parsing path, not the full search loop. Each proposal’s nearest neighbor among the eight shown members is found by minimum fragment-slot edit distance, calls that tie between multiple equally near members or resolve to no identifiable nearest member are excluded from the per-sample tally.

Table 8 reports all three samples. The three samples do not agree: the fraction of resolved calls nearest the falsely labeled-best member (the true worst) is 14.8%, 84.2%, and 46.2%, and the fraction of calls that tie or fail to resolve to a unique nearest member is also highly variable (13, 2, and 27 of 40) - plausibly reflecting how similar the eight randomly drawn archive members happen to be to each other in a given sample, which is not controlled across draws. The one pattern that holds in all three samples is that the genuinely best member is almost never the nearest neighbor regardless of its (false) label: 0, 3, and 0 of resolved calls, pooled 3/78 (3.8%), well below the 12.5% uniform base rate for an 8-member set. A single fixed-archive run of this probe therefore cannot support a specific claim about what proposals are tracking when the true and false quality signals conflict - the evidence here rules out “tracks true quality despite the false label” but does not establish a replicated alternative, seed 1’s 84.2% false-label-following, taken alone, would have suggested near-complete capture by the false label, while seed 0’s 14.8% would have suggested near-independence from either label, illustrating why this probe needs more than one fixed sample before its outcome is reported as a stable finding.

**Table 8:** Evidence-conflict probe across three independently drawn fixed archive samples, 40 calls each. “Other” denotes a resolved nearest neighbor that is neither the true-best nor the falsely labeled-best (true-worst) member.

| Sample seed | Resolved/40 | Nearest true-best | Nearest false-best | Nearest other | Tie/unresolved |
| --- | --- | --- | --- | --- | --- |
| 0 | 27 | 0 (0%) | 4 (14.8%) | 23 (85.2%) | 13 |
| 1 | 38 | 3 (7.9%) | 32 (84.2%) | 3 (7.9%) | 2 |
| 2 | 13 | 0 (0%) | 6 (46.2%) | 7 (53.8%) | 27 |
| Pooled | 78 | 3 (3.8%) | 42 (53.8%) | 33 (42.3%) | 42 |

The pooled row is reported for completeness but should not be read as a single stable rate: it averages over three samples whose individual false-label-following rates span a 69-percentage-point range, so the pooled 53.8% is an artifact of seed 1’s outlier sample rather than a representative summary. The pooled true-best rate (3.8%) is the only quantity that is consistent in direction across all three samples and is therefore the one load-bearing conclusion from this probe.

### K.6 Genuine reasoning-mode proposer

Ollama’s native think=true field is enabled for qwen3:8b, a mechanism distinct from the prompted v5_cot rationale, requesting JSON-formatted output simultaneously with think=true was found to silently suppress all reasoning output and is therefore not used, with JSON recovered instead by a hardened parser tolerant of markdown-fenced or prose-wrapped output. Under the default (unscored) prompt, four matched random-split seeds at *B* = 40 give Llama-comparable baselines (GA: 159.384, 112.501, 135.463, 100.518, random search: 139.085, 155.060, 166.663, 162.617) against reasoning-mode hypervolumes 101.998, 136.623, 93.679, and 89.201, giving median differences *−*26.55 (CI [*−*57.39, 24.12]) against GA and *−*55.04 (CI [*−*73.42*, −*18.44]) against random search. Diagnostics over 160 calls show 155 produced a reasoning trace (mean length 5,669 characters) and 0 mentioned potency or the objective. Under the scored prompt, all four seeds completed after raising Ollama’s per-call timeout to 300s (from the 120s default), following one timeout-related failure on the first attempt. Scored-prompt hypervolumes are 101.203, 98.645, 111.197, and 111.197 against the same GA and random-search baseline vectors as above, giving median differences *−*19.1 (GA) and *−*53.4 (random search), both CIs crossing zero. Diagnostics over 160 calls show 152 produced a reasoning trace (mean length 6,877 characters) and 124 (77.5%) mentioned potency, versus 0/155 under the default prompt - a sharp rise in measured engagement with the displayed evidence that does not translate into a hypervolume advantage over non-reasoning Llama 8B under the identical scored prompt (Table 4). Per-seed wall time for the default-prompt condition ranges from 2,801 to 5,290 seconds, roughly 5-10*×* the non-reasoning Llama or Qwen latency in Table 15.

### K.7 Prompt-variant latency

A dedicated timing run issues eight fresh Llama 3.1 8B calls per prompt variant against a fixed eight-member synthetic archive, measuring wall-clock seconds per call under identical local GPU conditions within one session. Mean seconds per call are 11.85 (default), 10.59 (v3_scored), 9.75 (v4_rank), 15.60 (v5_cot), and 13.99 (v6_shuffled). This fresh run’s default-prompt mean (11.85s) is higher than the 4.0s reported in Table 15, reflecting different GPU load/warm-state conditions between sessions rather than a changed model or prompt, absolute values are therefore not directly comparable across the two measurements, but the relative ordering within this single session is informative. The rationale-requiring v5_cot variant is the slowest, consistent with longer generated output and occasional rejection-retry overhead from the enforced non-empty rationale field, the shuffled and rank-only variants are within the same order of magnitude as the default and scored prompts, since none of these changes to the prompt content substantially change expected output length.

## L Structural inspection: additional detail

### L.1 3D rendering pipeline

Three-dimensional conformers for Figure 5 are generated from each best-found SMILES using RDKit’s ETKDG embedding followed by MMFF94 force-field geometry optimization (2,000 iterations) (Landrum and Contributors, 2026; Riniker and Landrum, 2015; Halgren, 1996), using the main project Python environment, and written to PDB files. These are rendered as ball-and-stick images using PyMOL’s util.cbaw coloring, a white background, and ray tracing (Schrödinger, LLC, 2026); PyMOL was installed via conda-forge in a separate, isolated conda environment created specifically for this rendering step. This isolation is deliberate: PyMOL’s dependency chain requires a newer numpy than the numpy version pinned throughout the rest of this project’s GP, active-learning, and statistical-testing code, and the rendering step does not need to share a Python environment with any numerical result reported elsewhere in this paper. No lipid geometry, charge state, or protonation beyond RDKit’s default valence-based structure generation is assumed, this is a visualization of real queried reagent identities, not a conformational or energetic claim about the physical lipid.

### L.2 Best-found lipids by method

For the objective-scored-prompt check’s seed 1, random split, Table 9 gives the best-found triple by frozen-oracle predicted HeLa+RAW264.7 sum for each method. Each is the arg-max over that method’s 40 queried candidates on this seed only, not a claim about the global optimum or about any other seed, different seeds and methods can and do find different best candidates.

**Table 9:**
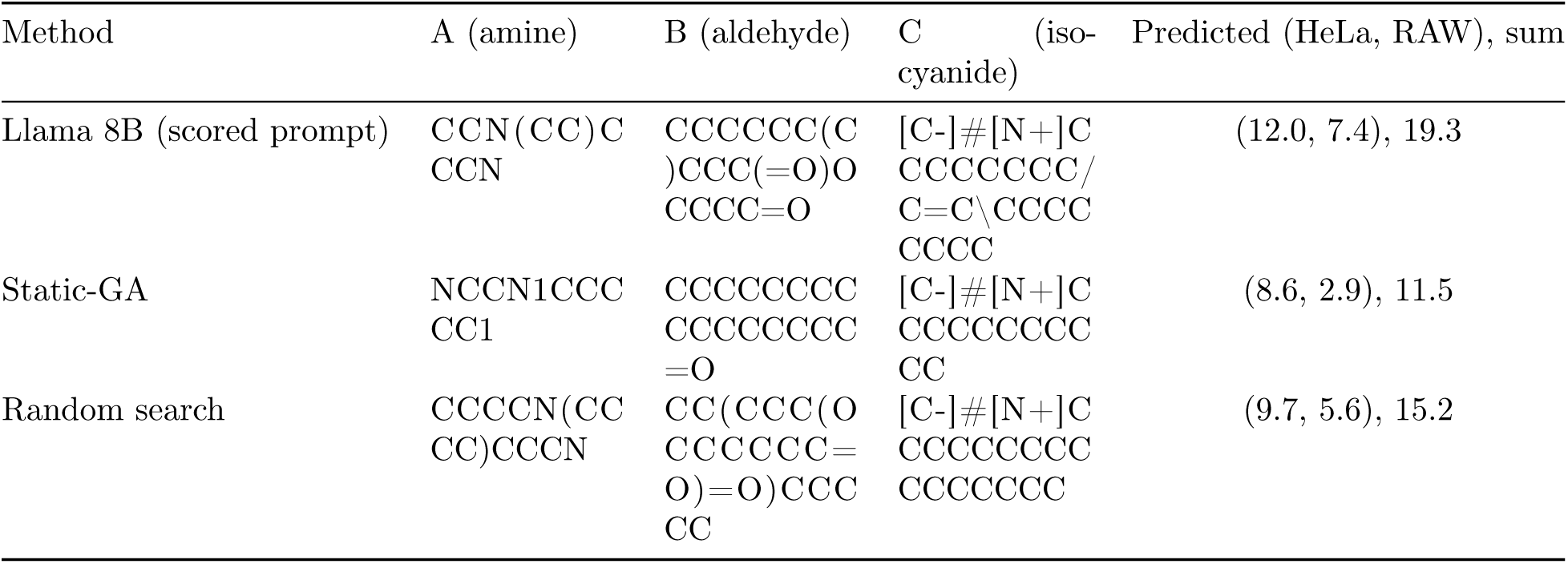
Best-found reagent triple per method, objective-scored-prompt check, random split, seed 1. SMILES strings wrap at arbitrary characters (seqsplit) since they contain no natural word breaks.

### L.3 Fragment-importance methodology and full range

For each of the 29 amine, 74 aldehyde, and 14 isocyanide values in the full extended vocabulary, every full-vocabulary candidate (30,044 total) containing that value is identified, and the frozen random-split *O*_GT_’s predicted HeLa+RAW264.7 sum is averaged across those candidates. Values appearing in fewer than 20 candidates are excluded to avoid small-sample means (this excludes none of the 29 amine or 14 isocyanide values, and a subset of the 74 aldehyde values). This is computed directly from the oracle’s posterior mean over the full combinatorial space - no search is run, no budget is charged, and no counterfactual pair or significance test is constructed, unlike the Hypothesis Ledger’s intended design (Appendix A).

Overall mean (across all 30,044 candidates) is 7.02 with standard deviation 1.41. The amine slot shows the widest range (3.20-9.61, a 6.4-point spread, dominated by cyclic tertiary amines with short linkers at the top and more flexible or doubly cyclic diamine-type structures at the bottom). The aldehyde slot’s range is 3.49-8.93 (5.4 points), the five lowest-scoring values are all doubly unsaturated (diene) long-chain tails, and the five highest are saturated, branched, shorter-chain linkers. The isocyanide slot’s range is narrow, 6.83-7.38 (0.55 points), and is dominated by tail length rather than unsaturation, though the two highest-scoring values are both mono-unsaturated. This means the apparent unsaturation association at slot C is both weaker in magnitude and opposite in direction to the one at slot B, so it should not be read as evidence for a general unsaturation-potency rule in this chemistry, any such claim would require an independently validated structure-activity study, not a correlational read of one fitted oracle.

## M Literature-lipid domain-coverage check

Three clinically deployed ionizable lipids (Escalona-Rayo et al., 2023) are checked against this paper’s real decompose_lipid routine (validated to exactly round-trip all 1,200 labeled AGILE lipids, Section 2.3): MC3 (DLin-MC3-DMA, PubChem CID 49785164), SM-102 (PubChem CID 126697616), and ALC-0315 (PubChem CID 122666778). Canonical SMILES for each were obtained from their respective PubChem records (Kim et al., 2023) and independently verified by computing each SMILES’s molecular formula with RDKit and matching it exactly against the formula listed on that compound’s own PubChem page (C43H79NO2, C44H87NO5, and C48H95NO5, respectively) before use, rather than trusting a search-engine summary.

All three SMILES parse validly in RDKit, but none match the Ugi-3CR core substructure. This is the expected outcome: MC3 is a dilinoleyl ester tertiary amine, while SM-102 and ALC-0315 are branched-ester tertiary amines with hydroxyl or ether linkers (Escalona-Rayo et al., 2023; Kim et al., 2023); none are products of an Ugi three-component reaction, so none should be expected to share this paper’s core scaffold. The check is reported because it was previously named in the main text as a planned but unreported comparison (Recall@*k* against MC3/SM-102/ALC-0315, Section 2.3), running it and disclosing the negative result removes that gap rather than leaving a stated plan silently unaddressed. No oracle potency prediction is computed for these three compounds, since the frozen *O*_GT_ requires a Morgan-fingerprint embedding of an assembled Ugi-3CR fragment triple and these molecules supply no such triple.

## N Panel interpretation

Figure 1 presents the budget-accounted architecture, Algorithm 1 and Section 2.1 define implemented budget accounting and endpoints. Hypothesis generation need not query the oracle, but counterfactual validation does. In the reported experiments, the Verifier is deterministic and the Ledger remains empty.

**Figure 1:**
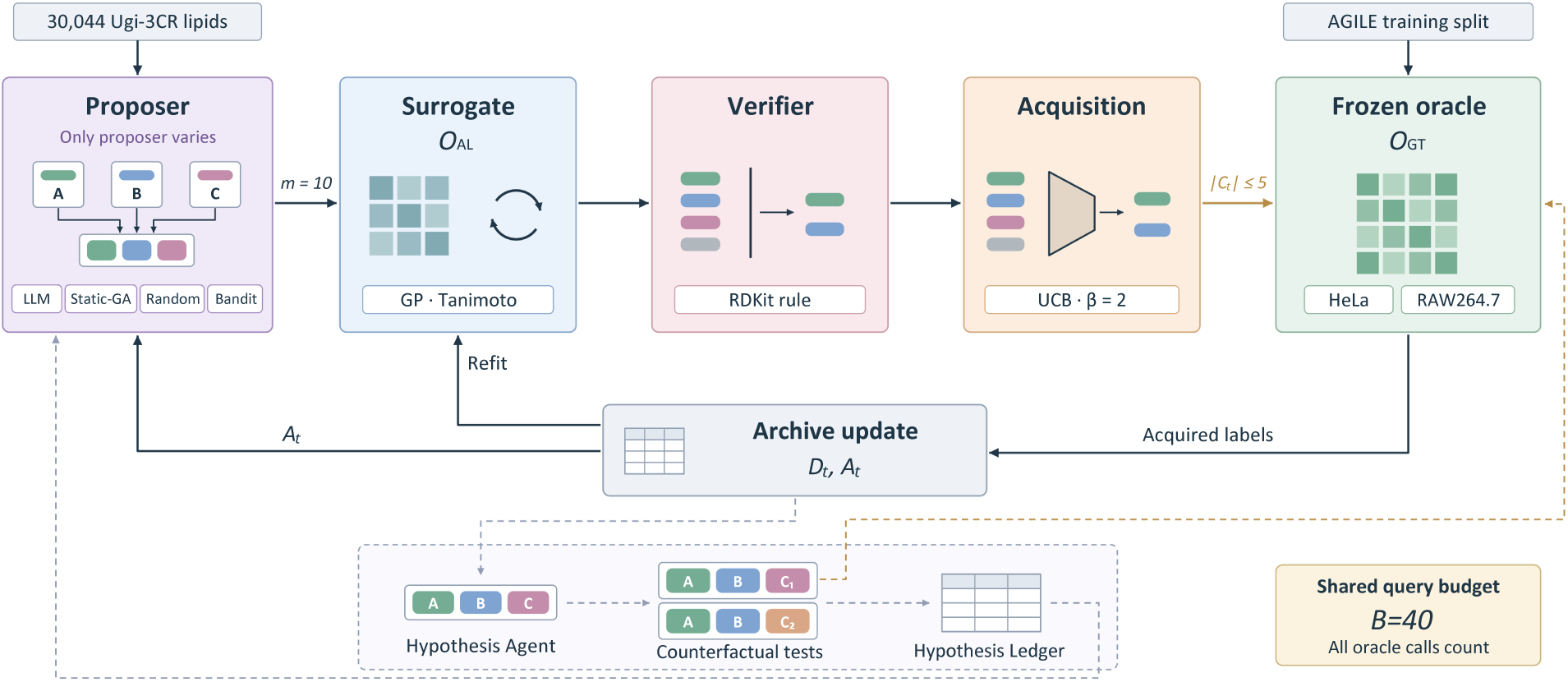
LEAF search loop under a shared oracle-query budget. Only the proposer varies across LLM, static-GA, random, and Thompson-bandit methods. Each round produces *m* = 10 reagent triples (*A, B, C*) from 30,044 Ugi-3CR lipids. The Tanimoto-GP surrogate *O*_AL_ scores candidates, an RDKit rule filters them, and UCB with *β* = 2 selects *|C_t_| ≤* 5 for evaluation by the frozen oracle *O*_GT_, fitted on the AGILE training split to predict HeLa and RAW264.7 potency. Acquired labels update *D_t_* and Pareto archive *A_t_*, supporting surrogate refitting and subsequent proposals. The dashed pathway depicts counterfactual hypothesis testing and Ledger feedback. Gold arrows denote oracle calls, including counterfactual evaluations, charged to the shared budget *B* = 40. Algorithm 1 specifies the accounting. The prompt-variant and reasoning-mode evidence-use checks (Section 3.4) run through this same budgeted loop with the Proposer box altered only in its input content or inference mode. The leakage probe and evidence-conflict probe are standalone diagnostics evaluated outside this loop and are not charged to *B*.

Figure 2 preserves held-out validity, median paired differences with bootstrap intervals, the zero reference line, unadjusted significance markers, and proposal diversity. Minimum edit distance counts changed reagent slots, from zero to three. Random search’s zero duplication is measured, not missing. Qwen 14B is the low-duplication exception among the four replicated models. Gemini’s aggregate duplication exceeds GA’s, but its chemotype-cold value of 19.2% is below GA’s reported 22.5%, split-specific results take precedence over blanket comparisons. Pilot and aggregate diagnostics remain distinct.

**Figure 2:**
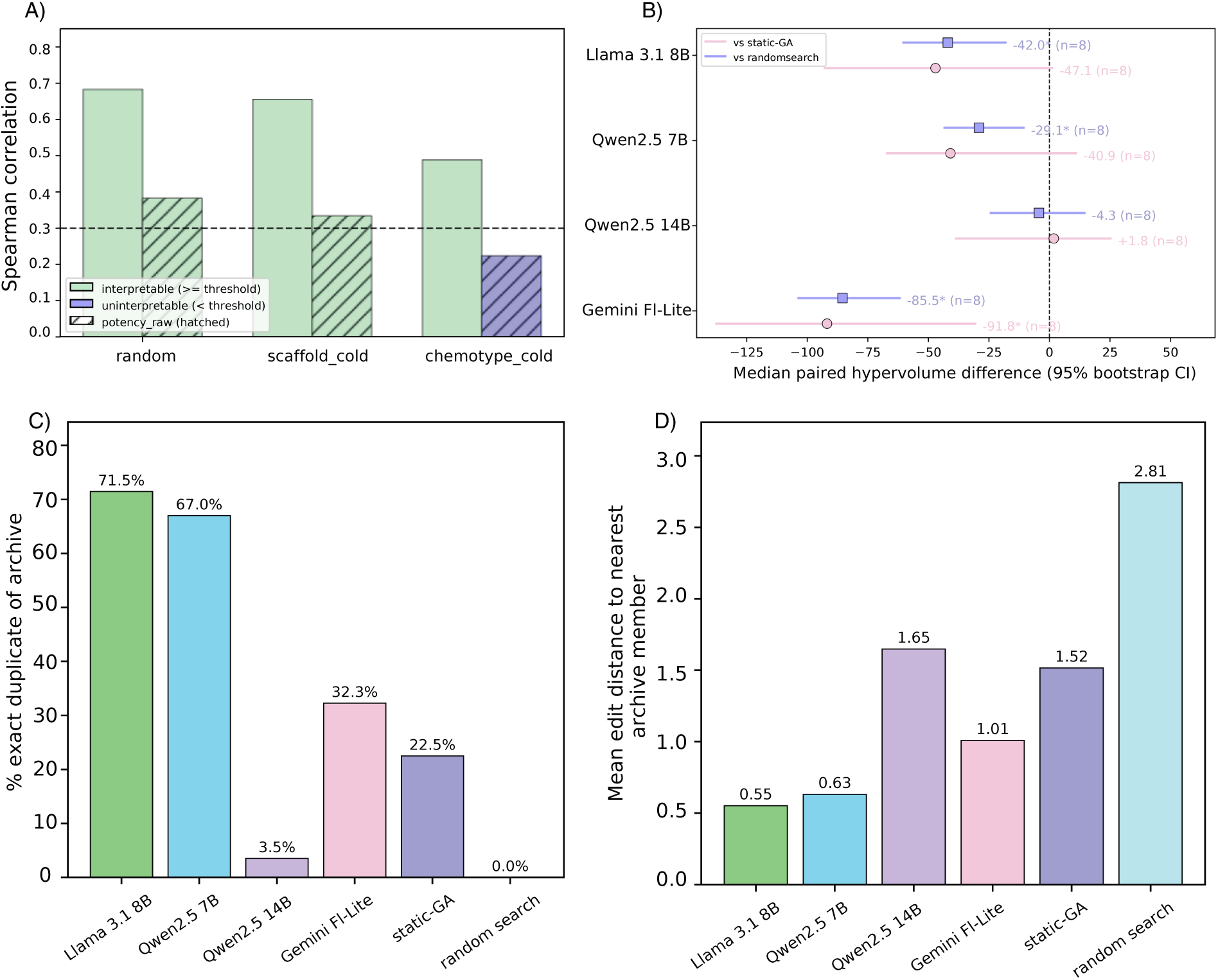
Oracle validity, proposal value, and diversity. **A:** Held-out potency correlations across three splits, the dashed line marks *ρ* = 0.3. Chemotype-cold RAW264.7 fails this gate. **B:** Random-split median paired hypervolume differences and 95% bootstrap intervals for four models, each with *n* = 8, asterisks mark unadjusted Wilcoxon *p <* 0.05, and the dashed line marks zero. The only positive estimate is Qwen 14B against static-GA, +1.8, not random search, *−*4.3, neither is significant. **C,D:** Exact archive-duplicate proposal rates and minimum fragment-slot edit distances from 0 to 3. Aggregate duplicate rates are 71.5%, 67.0%, 3.5%, and 32.3% for Llama 8B, Qwen 7B, Qwen 14B, and Gemini, versus 22.5% for GA and a measured 0% for random search. Corresponding distances are 0.55, 0.63, 1.65, and 1.01, versus 1.52 and 2.81. Table 13 separately reports the initial four-seed diagnostics. summaries only. Shared seeds and baseline runs invalidate independence-based pooled bootstrap intervals and Wilcoxon inferenc, no pooled inferential claim is made.

Figure 3 retains mean learning curves with standard deviations, early baseline gains and LLM flattening, log-scale nominal workload costs, and the non-significant 60-run duplication correlation. Its heatmaps include all 36 eight-seed comparisons, including the diversity prompt and non-LLM Thompson bandit. Groq’s three-seed experiment is excluded from those heatmaps, not treated as a complete eight-seed result.

**Figure 3:**
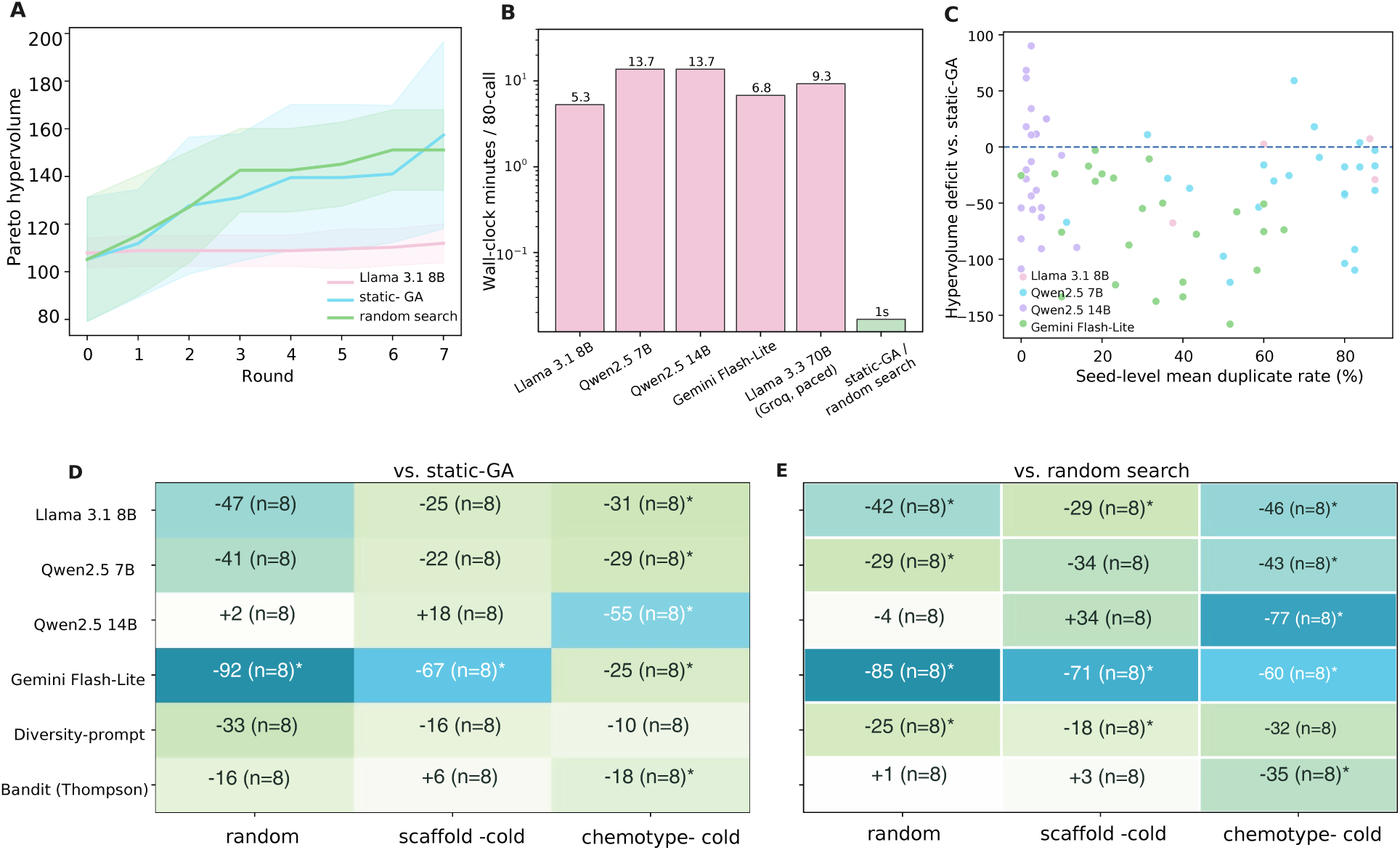
Learning dynamics, cost, and complete comparisons. **A:** Mean hypervolume with one standard deviation across eight random-split seeds, both baselines overtake Llama 8B within two to three rounds as its trajectory flattens. **B:** Log-scale nominal 80-call workload times, approximately 300-800 times the non-LLM cost, Table 15 gives timings. **C:** The 60-run robustness check finds no significant association between duplication and deficit, *ρ* = *−*0.12, *p* = 0.37. **D,E:** All 36 eight-seed comparisons across methods, splits, and baselines. Asterisks indicate unadjusted Wilcoxon *p <* 0.05. Positive cells are non-significant. The diversity prompt and non-LLM Thompson bandit are included.

**Figure 4:**
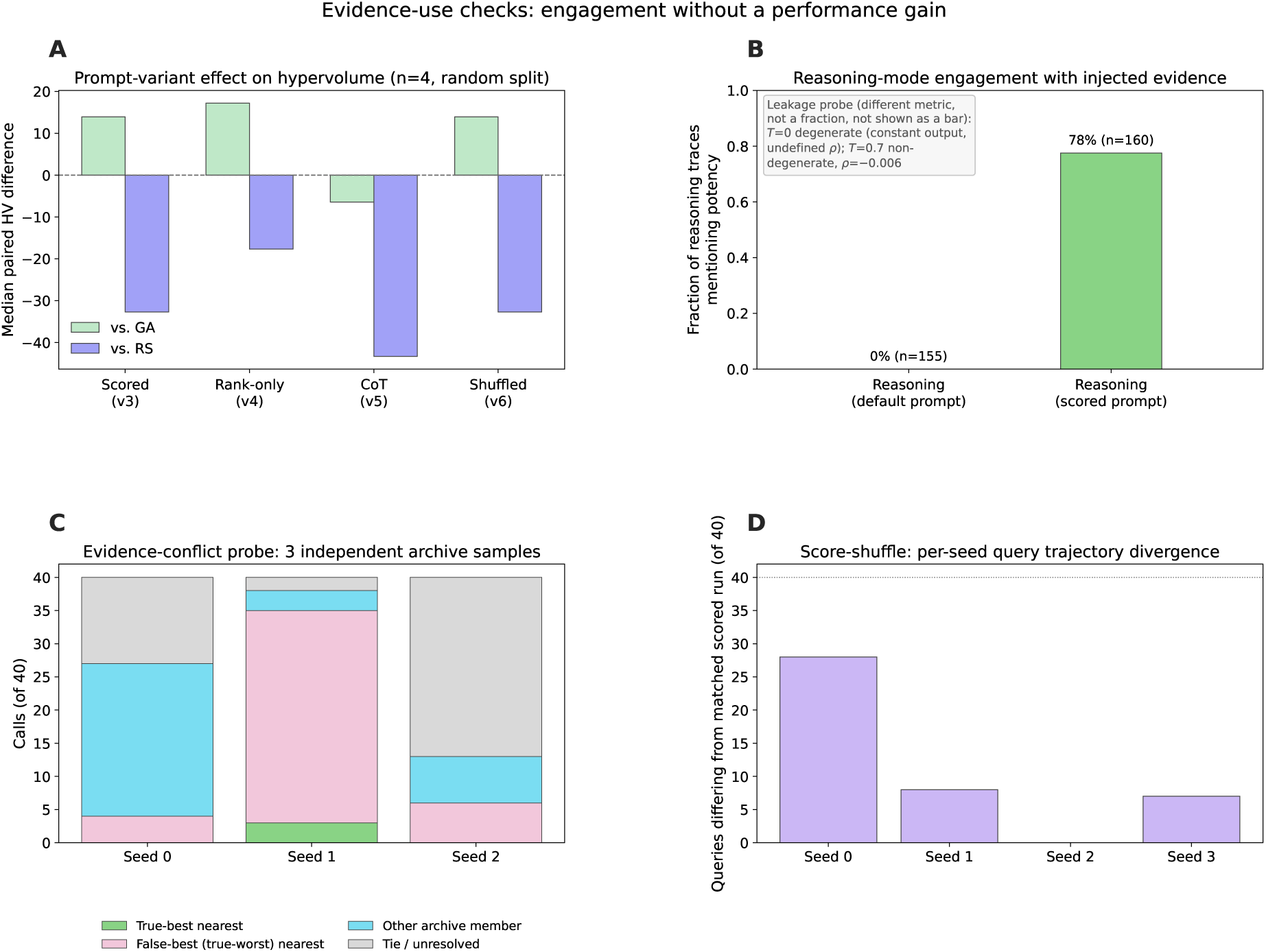
Evidence-use checks. **A:** Median paired hypervolume differences against GA and random search for each prompt variant (*n* = 4, random split). CIs cross zero in all cases. **B:** Fraction of reasoning-mode traces mentioning potency under the default versus scored prompt (0/155 versus 124/160). The leakage probe uses a different metric (Spearman correlation, degenerate at temperature zero and near-zero at temperature 0.7) and is annotated separately rather than plotted as a bar on this axis. **C:** Evidence-conflict probe nearest-neighbor outcome across three independently drawn fixed archive samples (40 calls each). The genuinely best member is almost never the nearest neighbor in any sample, but whether the falsely labeled-best member is followed instead varies from 15% to 84% across samples. The probe’s outcome is sample-dependent and not characterized by a single run. **D:** Per-seed count of queries differing from the matched correctly scored run under score-shuffle. Three of four seeds show real trajectory divergence despite similar aggregate hypervolume.

**Figure 5:**
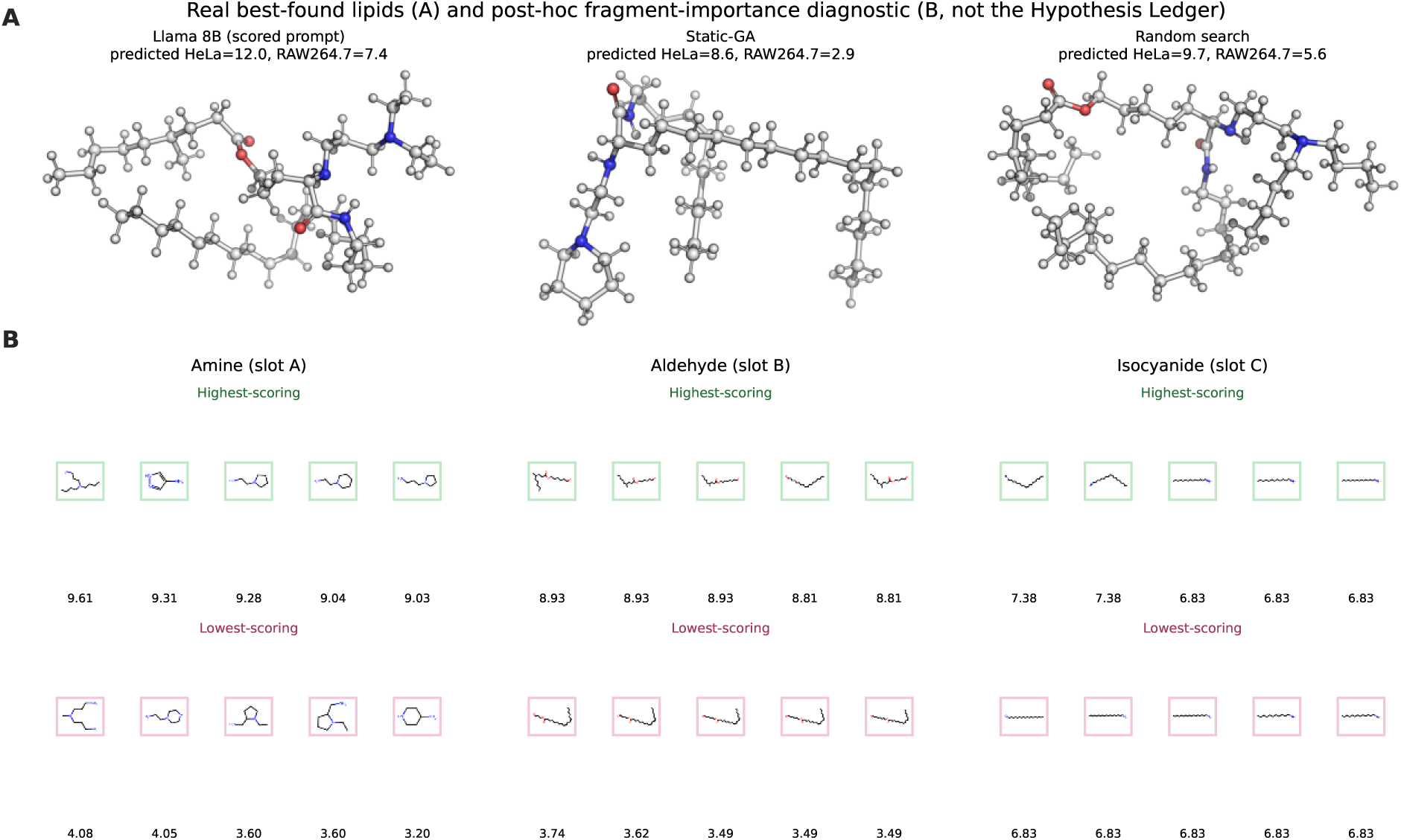
Real lipid structures and post-hoc fragment-importance diagnostic. **A:** 3D structure of each method’s best-found lipid (by frozen-oracle predicted HeLa+RAW264.7) among its 40 queried candidates, objective-scored-prompt check, random split, seed 1. A/B/C triples are the exact reagent identities queried under the shared budget. Conformers are generated with RDKit using ETKDG embedding and MMFF94 geometry optimization and rendered as ball-and-stick in PyMOL (Landrum and Contributors, 2026; Riniker and Landrum, 2015; Halgren, 1996; Schrödinger, LLC, 2026), run in an isolated environment so its dependencies do not affect the project’s pinned numerical stack (Appendix L). **B:** Post-hoc fragment-importance diagnostic over the full 30,044-candidate space under the frozen random-split oracle, independent of the (empty) Hypothesis Ledger: the five highest- and lowest-scoring fragment values at each reagent slot by mean predicted HeLa+RAW264.7 across every candidate containing that value, restricted to values appearing in at least 20 candidates. The opposite association with tail unsaturation at slots B and C indicates a slot-specific, non-generalizable relationship rather than a single interpretable structural rule.

## O Complete numerical tables

These eight tables retain every numerical entry in the original manuscript’s tables. Negative paired hypervolume differences favor the baseline. All tests are two-sided and unadjusted; eight seeds allow significance but do not guarantee statistical power. “GA” denotes static-GA and “RS” denotes random search.

**Table 10:**
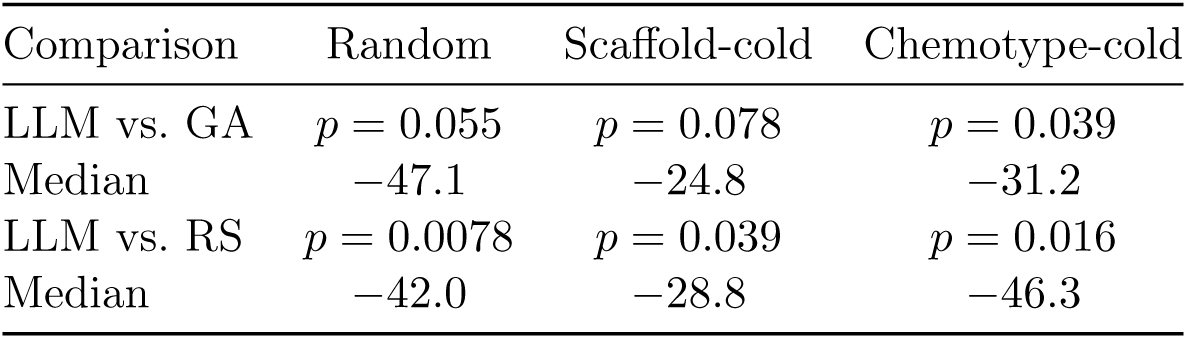
Llama 3.1 8B: paired hypervolume differences and Wilcoxon tests (*n* = 8).

| Comparison | Random | Scaffold-cold | Chemotype-cold |
| --- | --- | --- | --- |
| LLM vs. GA | $p = 0.055$ | $p = 0.078$ | $p = 0.039$ |
| Median | -47.1 | -24.8 | -31.2 |
| LLM vs. RS | $p = 0.0078$ | $p = 0.039$ | $p = 0.016$ |
| Median | -42.0 | -28.8 | -46.3 |

**Table 11:**
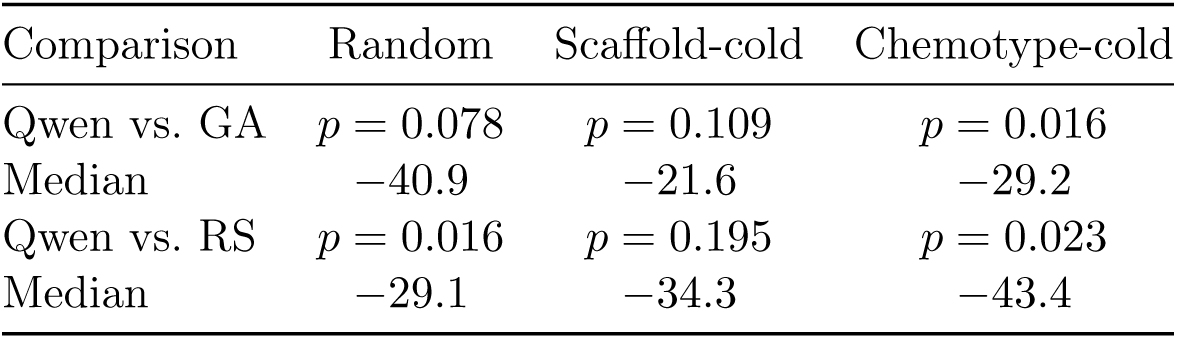
Qwen2.5 7B: paired hypervolume differences and Wilcoxon tests (*n* = 8).

| Comparison | Random | Scaffold-cold | Chemotype-cold |
| --- | --- | --- | --- |
| Qwen vs. GA | $p = 0.078$ | $p = 0.109$ | $p = 0.016$ |
| Median | -40.9 | -21.6 | -29.2 |
| Qwen vs. RS | $p = 0.016$ | $p = 0.195$ | $p = 0.023$ |
| Median | -29.1 | -34.3 | -43.4 |

**Table 12:** Qwen2.5 14B: paired hypervolume differences and Wilcoxon tests (*n* = 8).

| Comparison | Random | Scaffold-cold | Chemotype-cold |
| --- | --- | --- | --- |
| Qwen vs. GA | $p = 0.742$ | $p = 0.313$ | $p = 0.0078$ |
| Median | +1.8 | +18.3 | -55.1 |
| Qwen vs. RS | $p = 0.742$ | $p = 0.250$ | $p = 0.0078$ |
| Median | -4.3 | +33.9 | -77.0 |

**Table 13:** Initial proposal diagnostics (four seeds, one split). These are pool-proposal statistics, not directly measured fractions of selected oracle queries.

| Method | Crossover label | Archive duplicate | Edit distance |
| --- | --- | --- | --- |
| Llama 3.1 8B | 99.7% | 67.8% | 0.587 |
| Qwen2.5 7B | 56.2% | 62.5% | 0.650 |
| Static-GA | 0%<sup>†</sup> | 22.5% | 1.516 |
| Random search | 0%<sup>†</sup> | 0% | 2.812 |

<sup>†</sup>The non-LLM methods use different internal operator labels (e.g., `ga_crossover_mutation`); zero does not mean zero recombination.

**Table 14:** Matched de-duplication check: Llama 3.1 8B, random split, *B* = 15, *n* = 4. All 95% CIs cross zero.

| Comparison | No de-duplication | With de-duplication |
| --- | --- | --- |
| LLM vs. static-GA | Median $-17.6$ | Median $-4.7$ |
| LLM vs. random search | Median $-15.1$ | Median $+2.3$ |

**Table 15:**
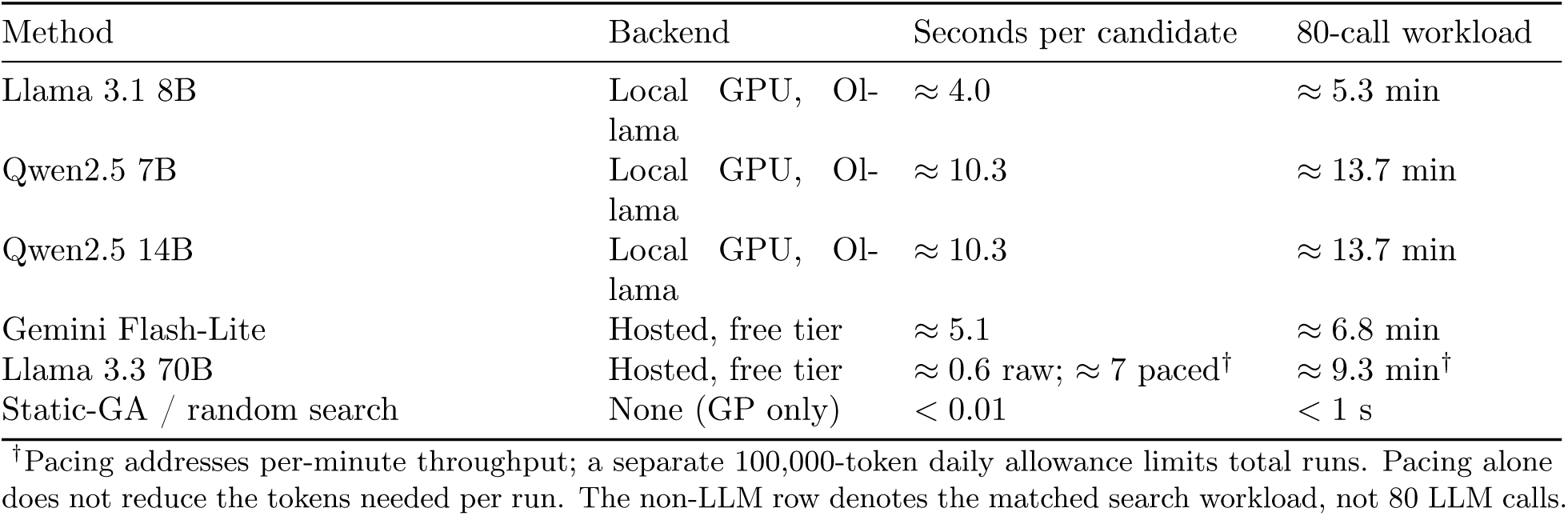
Measured candidate latency and 80-call workload cost. Times reproduce the original table; hosted retries, quota delays, and multi-day accumulation are additional.

**Table 16:**
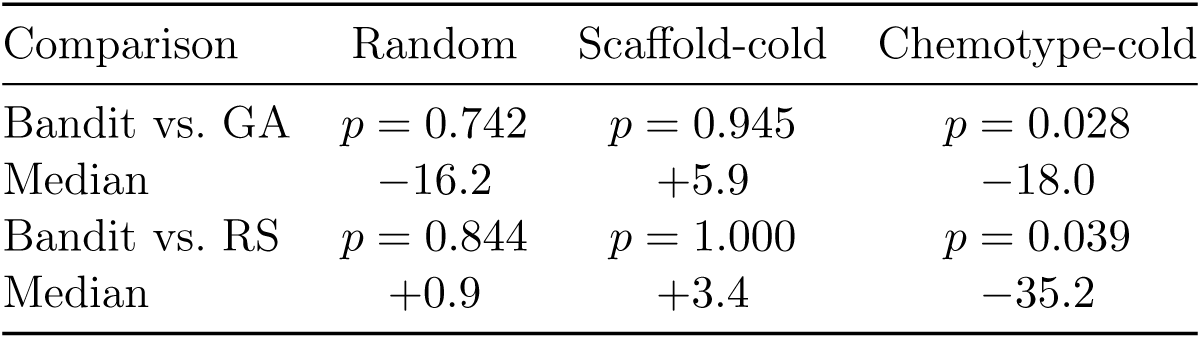
Thompson-sampling bandit: paired hypervolume differences and Wilcoxon tests (*n* = 8).

| Comparison | Random | Scaffold-cold | Chemotype-cold |
| --- | --- | --- | --- |
| Bandit vs. GA | $p = 0.742$ | $p = 0.945$ | $p = 0.028$ |
| Median | $-16.2$ | $+5.9$ | $-18.0$ |
| Bandit vs. RS | $p = 0.844$ | $p = 1.000$ | $p = 0.039$ |
| Median | $+0.9$ | $+3.4$ | $-35.2$ |

**Table 17:** Gemini Flash-Lite: paired hypervolume differences and Wilcoxon tests (*n* = 8).

| Comparison | Random | Scaffold-cold | Chemotype-cold |
| --- | --- | --- | --- |
| Gemini vs. GA | $p = 0.0078$ | $p = 0.0078$ | $p = 0.0078$ |
| Median | $-91.8$ | $-66.6$ | $-24.8$ |
| Gemini vs. RS | $p = 0.0078$ | $p = 0.0078$ | $p = 0.0078$ |
| Median | $-85.5$ | $-71.1$ | $-60.2$ |

